## Supplementary Material - Equation and Graphics for "A Bayesian Neural Ordinary Differential Equations Framework to Study the Effects of Chemical Mixtures on Survival"

March 14, 2025

Virgile BAUDROT[1*],Nina CEDERGREEN[2],Thomas KLEIBER[1],André GERGS[3], Sandrine CHARLES[4]

**1** Qonfluens, Montpellier, France

**2** University of Copenhagen, Copenhagen, Denmark

**3** Bayer Crop Sciences, Mannheim, Germany

**4** Université Claude Bernard Lyon 1, Villeurbanne, France

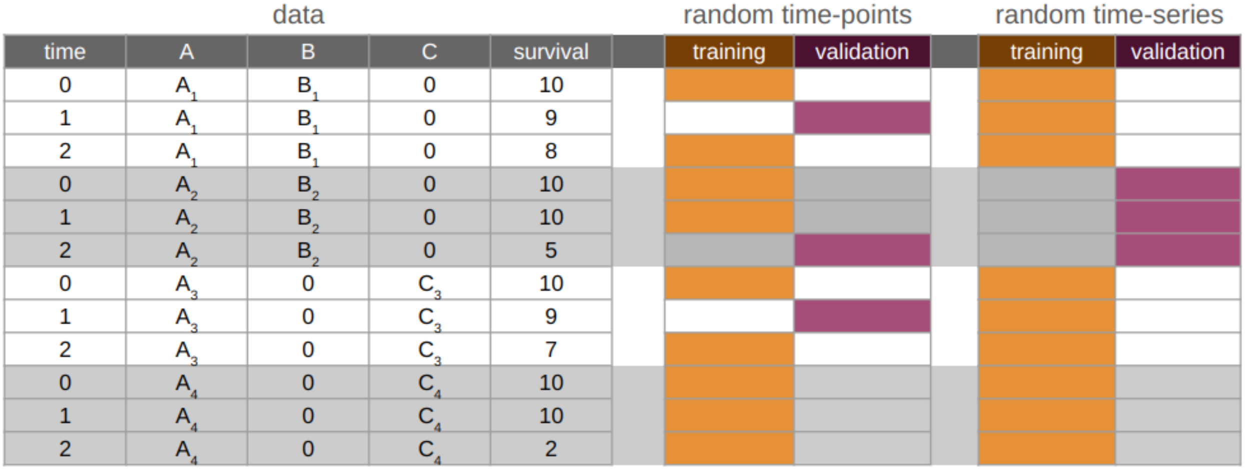


Figure 1: Visualization of dataset splitting for two different binary mixtures (AB and AC), both sharing compound A. In orange, the training subset is shown, where the model is fitted, and parameters are estimated. In purple, the validation subset is displayed, where model predictions are compared for validation. The "random time-points" columns represent a random selection of 90% of time points across the entire dataset for calibration, with the remaining 10% used for validation. The "random time-series" columns depict the selection of 90% of time series for calibration and 10% for validation, respectively.

| | time | A | B | C | survival | training | validation |
|---|---|---|---|---|---|---|---|
| AB | 0 | $A_1$ | $B_1$ | 0 | 10 | | |
| | 1 | $A_1$ | $B_1$ | 0 | 9 | | |
| | 2 | $A_1$ | $B_1$ | 0 | 8 | | |
| | 0 | $A_2$ | $B_2$ | 0 | 10 | | |
| | 1 | $A_2$ | $B_2$ | 0 | 10 | | |
| | 2 | $A_2$ | $B_2$ | 0 | 5 | | |
| AC | 0 | $A_3$ | 0 | $C_3$ | 10 | | |
| | 1 | $A_3$ | 0 | $C_3$ | 9 | | |
| | 2 | $A_3$ | 0 | $C_3$ | 7 | | |
| | 0 | $A_4$ | 0 | $C_4$ | 10 | | |
| | 1 | $A_4$ | 0 | $C_4$ | 10 | | |
| | 2 | $A_4$ | 0 | $C_4$ | 2 | | |
| BC | 0 | 0 | $B_5$ | $C_5$ | 10 | | |
| | 1 | 0 | $B_5$ | $C_5$ | 5 | | |
| | 2 | 0 | $B_5$ | $C_5$ | 1 | | |
| | 0 | 0 | $B_6$ | $C_6$ | 10 | | |
| | 1 | 0 | $B_6$ | $C_6$ | 9 | | |
| | 2 | 0 | $B_6$ | $C_6$ | 3 | | |

Figure 2: Illustration of the selection scheme used for validation on uncalibrated mixtures. Across the entire dataset, all mixtures containing a specific pair of active ingredients (e.g., $AB$) are excluded from the calibration dataset and reserved exclusively for validation.

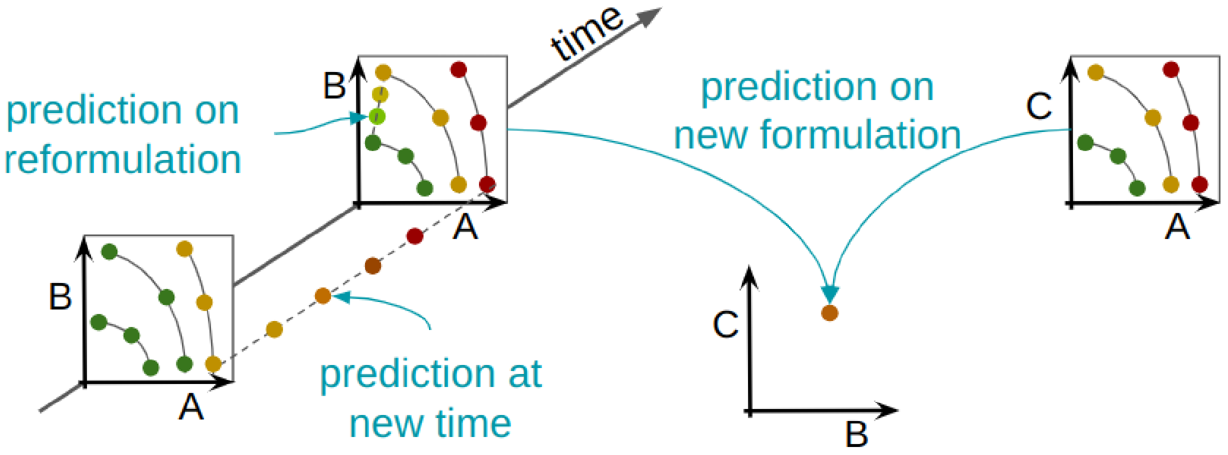


Figure 3: The figure illustrates three prediction types: (1) predictions at a new time point with the identical mixture; (2) reformulation predictions (same active ingredients but in different proportions); and (3) predictions for not yet tested newly composed mixtures of tested active ingredients.

### 1 Summary of the four datasets

#### 1.1 Dataset 1

Dataset 1 is made of 244 time-series covering 21 mixtures, given 1482 survival data points.

The active ingredients are: bixafen, fluopyram, prothioconazole, spiroxamine, tebuconazole and trifloxystrobin.

And the 21 mixtures are given as follows:

| Mixtures | Number of time series |
|---|---|
| bixafen | 13 |
| bixafen, fluopyram | 6 |
| bixafen, fluopyram, prothioconazole | 6 |
| bixafen, prothioconazole | 6 |
| bixafen, prothioconazole, spiroxamine | 6 |
| bixafen, prothioconazole, tebuconazole | 6 |
| bixafen, tebuconazole | 6 |
| fluopyram | 6 |
| fluopyram, spiroxamine | 6 |
| fluopyram, tebuconazole | 6 |
| fluopyram, trifloxystrobin | 18 |
| prothioconazole | 25 |
| prothioconazole, spiroxamine | 6 |
| prothioconazole, spiroxamine, tebuconazole | 6 |
| prothioconazole, spiroxamine, trifloxystrobin | 6 |
| prothioconazole, tebuconazole | 18 |
| prothioconazole, trifloxystrobin | 12 |
| spiroxamine | 22 |
| tebuconazole | 22 |
| tebuconazole, trifloxystrobin | 12 |
| trifloxystrobin | 30 |

Table 1: Dataset 1 mixtures.

| Mixtures | Pairwise availability of data |
|---|---|
| bixafen, fluopyram | data |
| bixafen, prothioconazole | data |
| bixafen, spiroxamine | no data |
| bixafen, tebuconazole | data |
| bixafen, trifloxystrobin | no data |
| fluopyram, prothioconazole | no data |
| fluopyram, spiroxamine | data |
| fluopyram, tebuconazole | no data |
| fluopyram, trifloxystrobin | data |
| prothioconazole, spiroxamine | data |
| prothioconazole, tebuconazole | data |
| prothioconazole, trifloxystrobin | data |
| spiroxamine, tebuconazole | no data |
| spiroxamine, trifloxystrobin | no data |
| tebuconazole, trifloxystrobin | data |

Table 2: Pairwise for dataset 1.

#### 1.2 Dataset 2

Dataset 2 is made of 120 time-series covering 9 mixtures, given 662 survival data points.

The active ingredients are: deltamethrin, flupyradifurone, spiromesifen, tralomethrin and triazophos.

| Mixtures | Number of time series |
|---|---|
| deltamethrin | 39 |
| deltamethrin, flupyradifurone | 12 |
| deltamethrin, tralomethrin | 12 |
| deltamethrin, triazophos | 8 |
| flupyradifurone | 7 |
| flupyradifurone, spiromesifen | 7 |
| spiromesifen | 13 |
| tralomethrin | 14 |
| triazophos | 8 |

Table 3: Dataset 2 mixture.

#### 1.3 Dataset 3

Dataset 3 is made of 130 time-series covering 11 mixtures, given 727 survival data points.

The active ingredients are: beta-cyfluthrin, clothianidin, cyfluthrin, imidacloprid, thiacloprid and thiodicarb.

#### 1.4 Dataset 4

Dataset 4 is made of 126 time-series covering 12 mixtures, given 758 survival data points.

The active ingrediens are: aclonifen, diflufenican, flufenacet, flurtamone and metribuzin.

| Mixture | Pairwise availability of data |
|---|---|
| deltamethrin, flupyradifurone | data |
| deltamethrin, spiromesifen | no data |
| deltamethrin, tralomethrin | data |
| deltamethrin, triazophos | data |
| flupyradifurone, spiromesifen | data |
| flupyradifurone, tralomethrin | no data |
| flupyradifurone, triazophos | no data |
| spiromesifen, tralomethrin | no data |
| spiromesifen, triazophos | no data |
| tralomethrin, triazophos | no data |

Table 4: Pairwise for dataset 2.

| Mixtures | Number of time series |
|---|---|
| beta-cyfluthrin | 25 |
| beta-cyfluthrin, clothianidin | 6 |
| beta-cyfluthrin, thiacloprid | 6 |
| clothianidin | 7 |
| clothianidin, imidacloprid, thiodicarb | 6 |
| cyfluthrin | 13 |
| cyfluthrin, imidacloprid | 7 |
| imidacloprid | 21 |
| imidacloprid, thiodicarb | 19 |
| thiacloprid | 7 |
| thiodicarb | 13 |

Table 5: Dataset 3 mixture.

| Mixtures | Pairwise availability of data |
|---|---|
| beta-cyfluthrin, clothianidin | data |
| beta-cyfluthrin, cyfluthrin | no data |
| beta-cyfluthrin, imidacloprid | no data |
| beta-cyfluthrin, thiacloprid | data |
| beta-cyfluthrin, thiodicarb | no data |
| clothianidin, cyfluthrin | no data |
| clothianidin, imidacloprid | no data |
| clothianidin, thiacloprid | no data |
| clothianidin, thiodicarb | no data |
| cyfluthrin, imidacloprid | data |
| cyfluthrin, thiacloprid | no data |
| cyfluthrin, thiodicarb | no data |
| imidacloprid, thiacloprid | no data |
| imidacloprid, thiodicarb | data |
| thiacloprid, thiodicarb | no data |

Table 6: Pairwise for dataset 3.

##### 1.5 Checking potential effect of co-formulants

In order to make sure that co-formulant do not have an effect on the survival, we compared the LC50 predicted with the formulation with the one predicted with active ingredients. We did it on single compound formulation which share the same co-formulant are those used in formulation with mixture of several active ingredients. No additional effects of co-formulant were detected.

In practice, using for instance a time series of a study on spiroxamine formulation at 97.8. One one side, we fit the data considering the formulation concentration and then compute the LC50 (which was 17.516 [15.817, 18.245]) and one the other side we fit data considering the active ingredient concentration only, where we obtain the LC50 (17.121 [15.541, 17.844]). As a measure of discrepancy, we compared the ratio between the two LC50 with the ratio of active ingredients in the formulation. In this example of spiroxamine, it corresponds to LC50 active ingredient / LC 50 formulation = 17.121 / 17.516 with 97.8 ), and the difference gives -0.00055, leading to consider no effect of the co-formulant in this situation. We did the same procedure for all sets of time series with a single active ingredient, and the results are given in the graphics here-after. This difference

| Mixtures | Number of time series |
|---|---|
| aclonifen | 25 |
| aclonifen, diflufenican | 6 |
| aclonifen, diflufenican, flufenacet | 7 |
| aclonifen, flufenacet | 6 |
| aclonifen, flurtamone | 6 |
| diflufenican | 12 |
| diflufenican, flufenacet | 14 |
| diflufenican, flufenacet, flurtamone | 6 |
| diflufenican, metribuzin | 6 |
| flufenacet | 21 |
| flurtamone | 7 |
| metribuzin | 10 |

Table 7: Dataset 4 mixture.

| Mixtures | Pairwise availability of data |
|---|---|
| aclonifen diflufenican | data |
| aclonifen flufenacet | data |
| aclonifen flurtamone | data |
| aclonifen metribuzin | no data |
| diflufenican flufenacet | data |
| diflufenican flurtamone | no data |
| diflufenican metribuzin | data |
| flufenacet flurtamone | no data |
| flufenacet metribuzin | no data |
| flurtamone metribuzin | no data |

Table 8: Pairwise for dataset 4.

never exceeds 0.015.

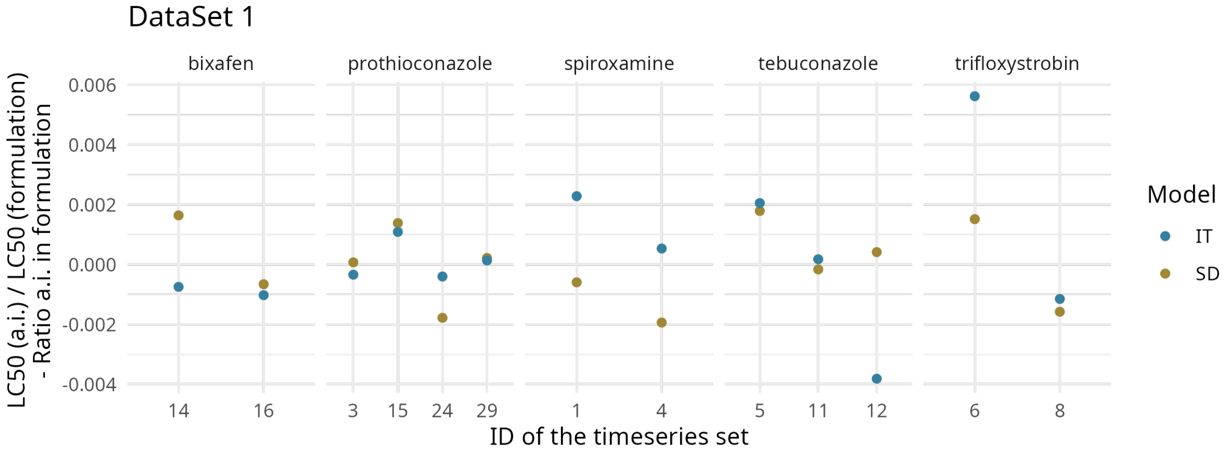

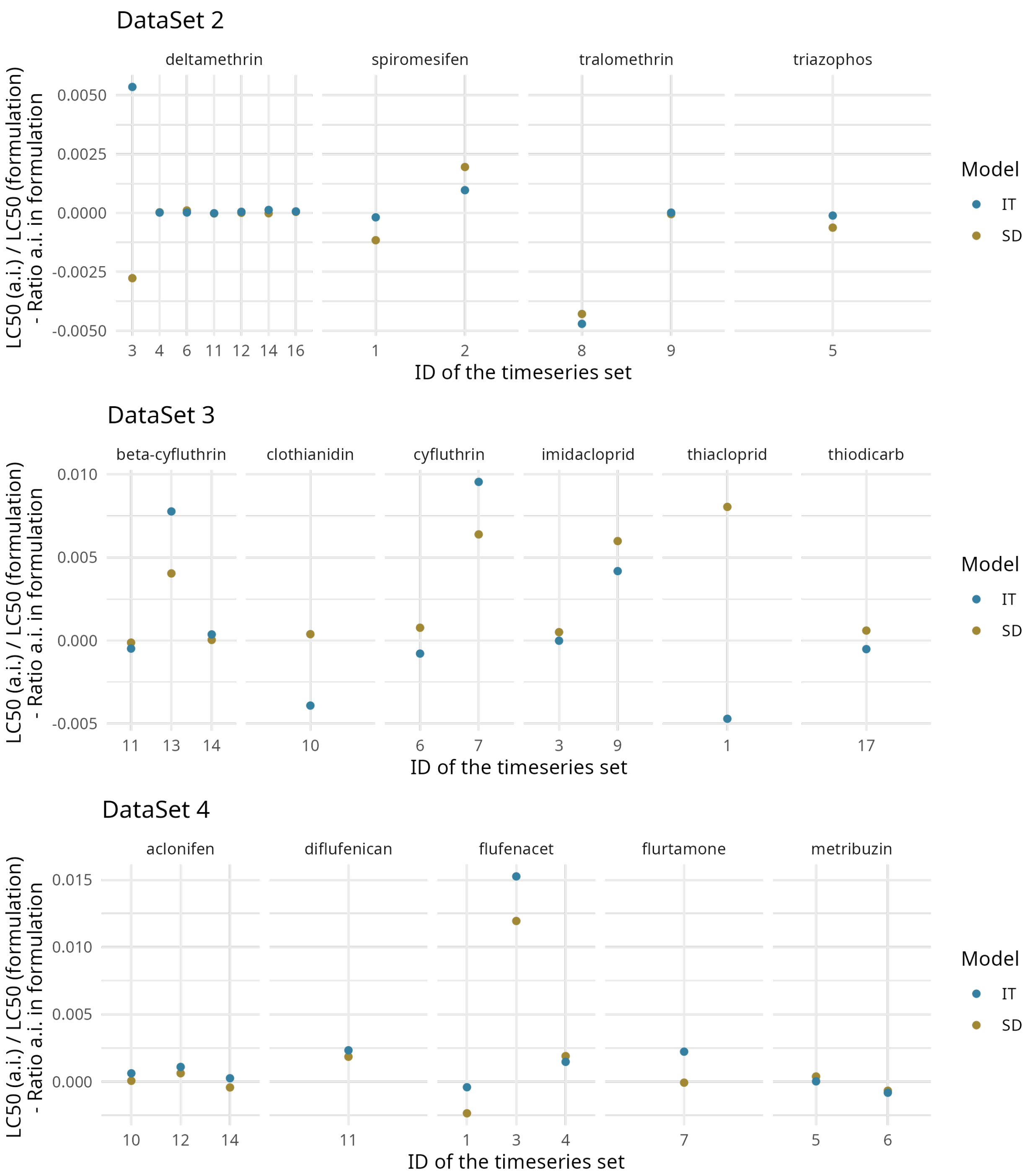

DataSet 2
deltamethrin
spiromesifen
tralomethrin
triazophos
LC50 (a.i.) / LC50 (formulation)
- Ratio a.i. in formulation
ID of the timeseries set
Model
IT
SD
DataSet 3
beta-cyfluthrin
clothianidin
cyfluthrin
imidacloprid
thiacloprid
thiodicarb
DataSet 4
aclonifen
diflufenican
flufenacet
flurtamone
metribuzin

#### 1.6 Goodness of Fit of Cross-Validation

To assess the quality of the calibration output, we used the model performance criteria recommended by EFSA in its Scientific Opinion on TKTD modeling efsa2018tktd, including the Posterior Prediction Check (PPC) and the the Normalized Root Mean Square Error (NRMSE).

The PPC enables a comparison between the predicted and observed survival counts, summarizing the predictions by their mean and the 95% uncertainty range. The percentage of predictions that capture the observed data serves as a quantitative goodness-of-fit criterion.

The NRMSE is defined by the equation:

$$\text{NRMSE} = \frac{1}{\frac{1}{N}\sum_i y_{obs,i}} \sqrt{\frac{1}{N}\sum_i \left(y_{obs,i} - y_{pred,i}\right)^2} \tag{1}$$

where $y_{obs,i}$ and $y_{pred,i}$, $i = 1, N$, are the observed and predicted values, respectively, across $N$ data points.

Since Bayesian inference was applied, we included the Widely Applicable Information Criterion (WAIC) to estimate the effective number of parameters, accounting for overfitting watanabe2013widely. The WAIC is defined as:

$$WAIC = -2\,(\text{lpd} - \text{pWAIC}) = -2\left(\sum_i \log(E(p(y|\theta))) - \sum_i \text{var}(\log(p(y|\theta))\right) \tag{2}$$

where lpd is the log-point-wise predictive density and pWAIC represents the effective number of parameters.

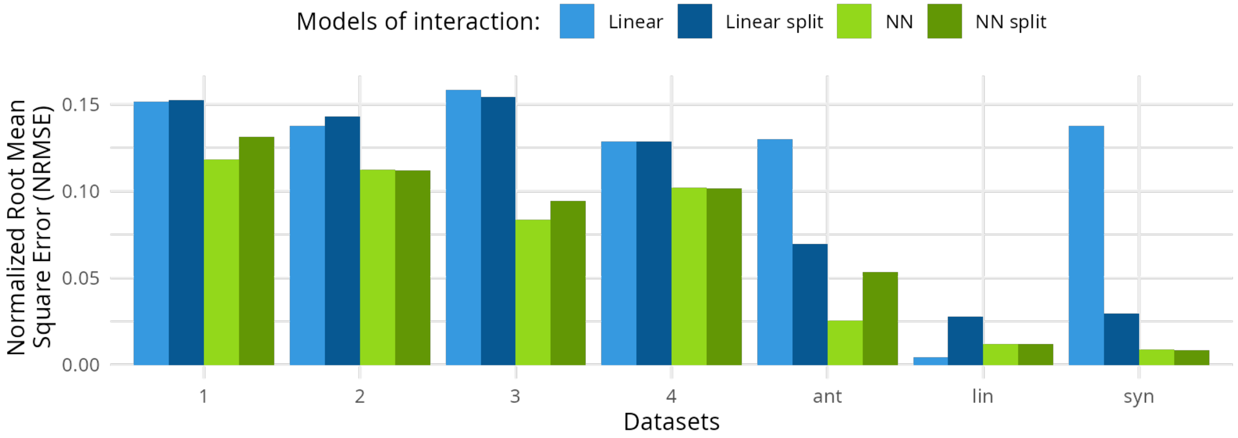

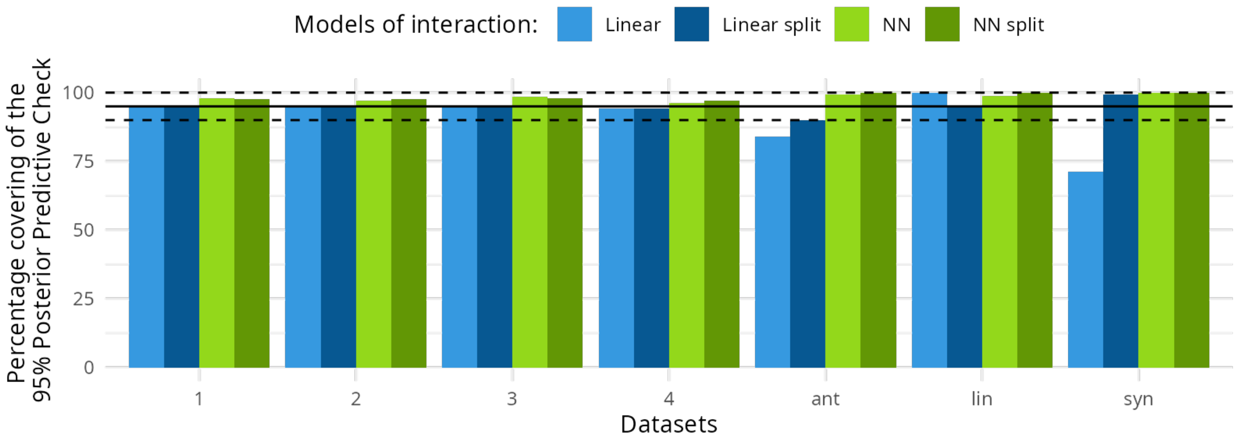

Models of interaction:
Linear
Linear split
NN
NN split
Percentage covering of the 95% Posterior Predictive Check
100
75
50
25
0
1
2
3
4
ant
lin
syn
Datasets


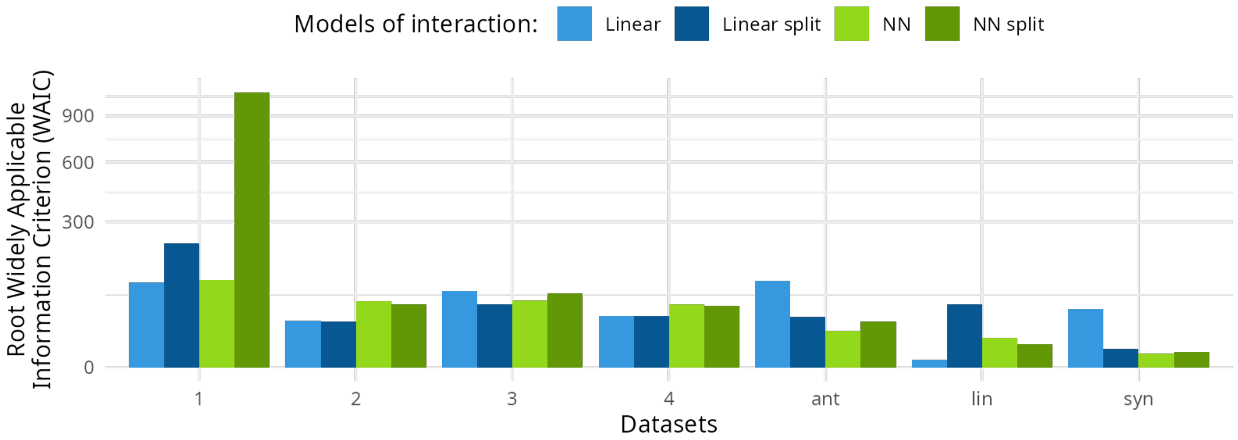

Models of interaction:
Linear
Linear split
NN
NN split
Root Widely Applicable Information Criterion (WAIC)
900
600
300
0
1
2
3
4
ant
lin
syn
Datasets

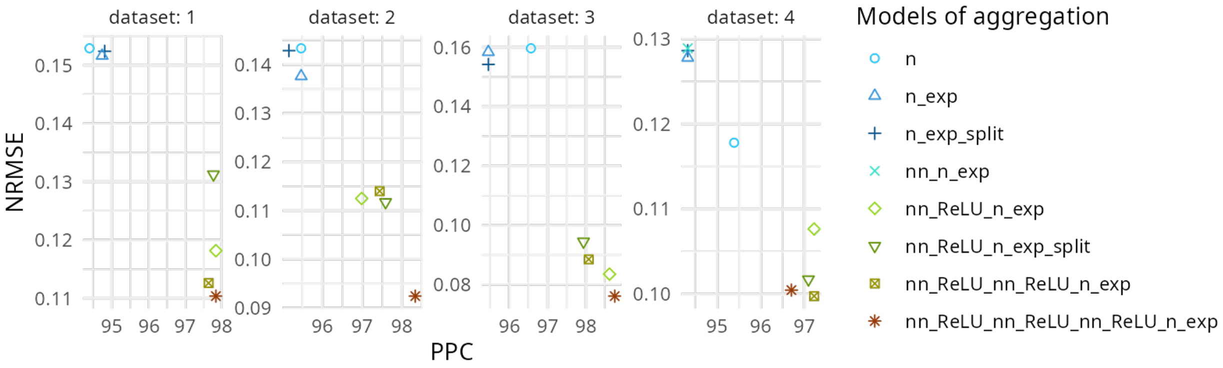


Figure 4: Scatter plots illustrate the quality of calibration results for three artificial data sets fitted to five generic models (upper panel) and for four experimental data sets fitted to eight different models (lower panel). The covering of data by the Posterior Predictive Check (PPC) ($x$-axis) and NRMSE ($y$-axis) serve as goodness-of-fit metrics, with lower values indicating higher fit quality. In the "Models of aggregations" legend, `n` stands for the simplest linear model with $n$ compounds $(b+\sum_{i=1}^{n} a_i C_i)$; `n_exp` for the linear model with an exponential transformation $(\exp(b+\sum_{i=1}^{n} a_i C_i))$; `n_exp_split` is the `n_exp` model with the splitting of $\alpha$; `nn_n_exp` is the application of a perceptron matrix of size $n \times n$ without activation function before the `n_exp` model; and `nn_ReLU_n_exp` is the application of an $n \times n$ layer with a `ReLU` function before the `n_exp` model. The remaining models with the `nn_ReLU` prefix involve a sequence of perceptron layers before the `n_exp` model. See Manuscript for details. The data sets called "add", "ant" and "syn" were simulated under additive, antagonism and synergism interaction hypotheses, respectively.

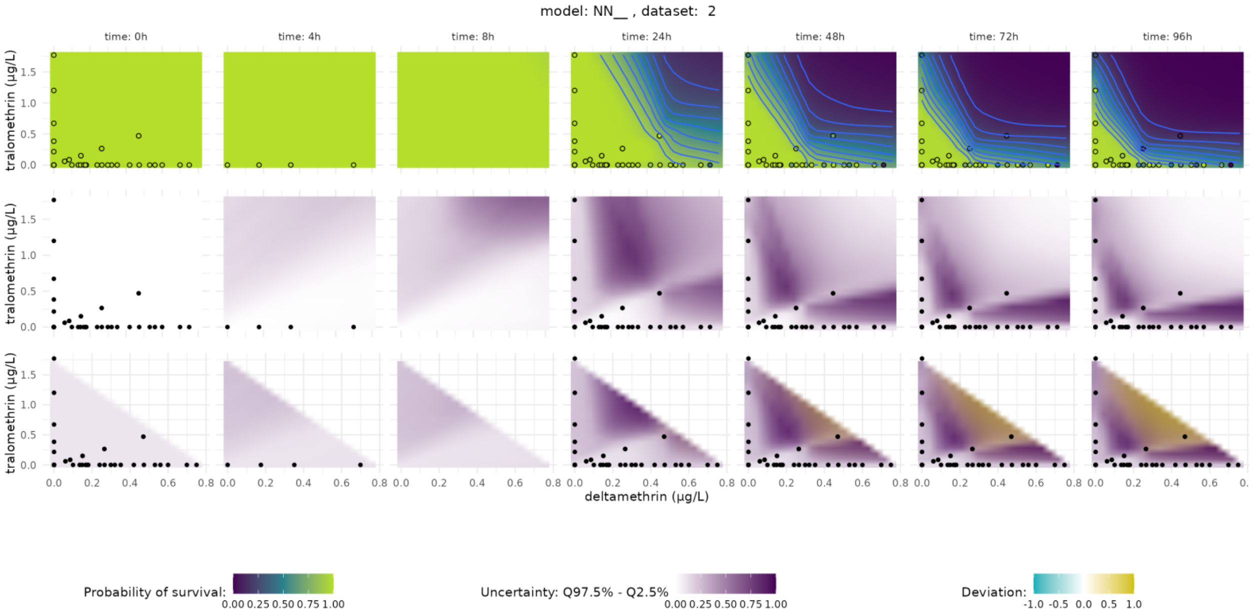

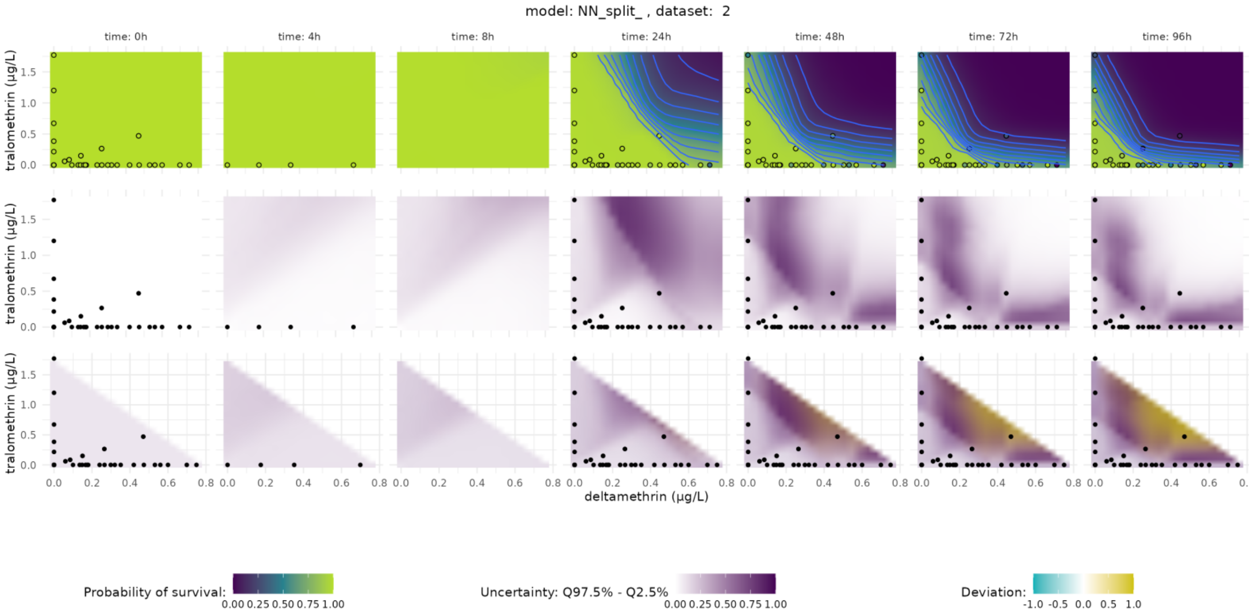
model: NN_split_ , dataset: 2
time: 0h
time: 4h
time: 8h
time: 24h
time: 48h
time: 72h
time: 96h
tralomethrin (µg/L)
deltamethrin (µg/L)
Probability of survival:
0.00 0.25 0.50 0.75 1.00
Uncertainty: Q97.5% - Q2.5%
0.00 0.25 0.50 0.75 1.00
Deviation:
-1.0 -0.5 0.0 0.5 1.0

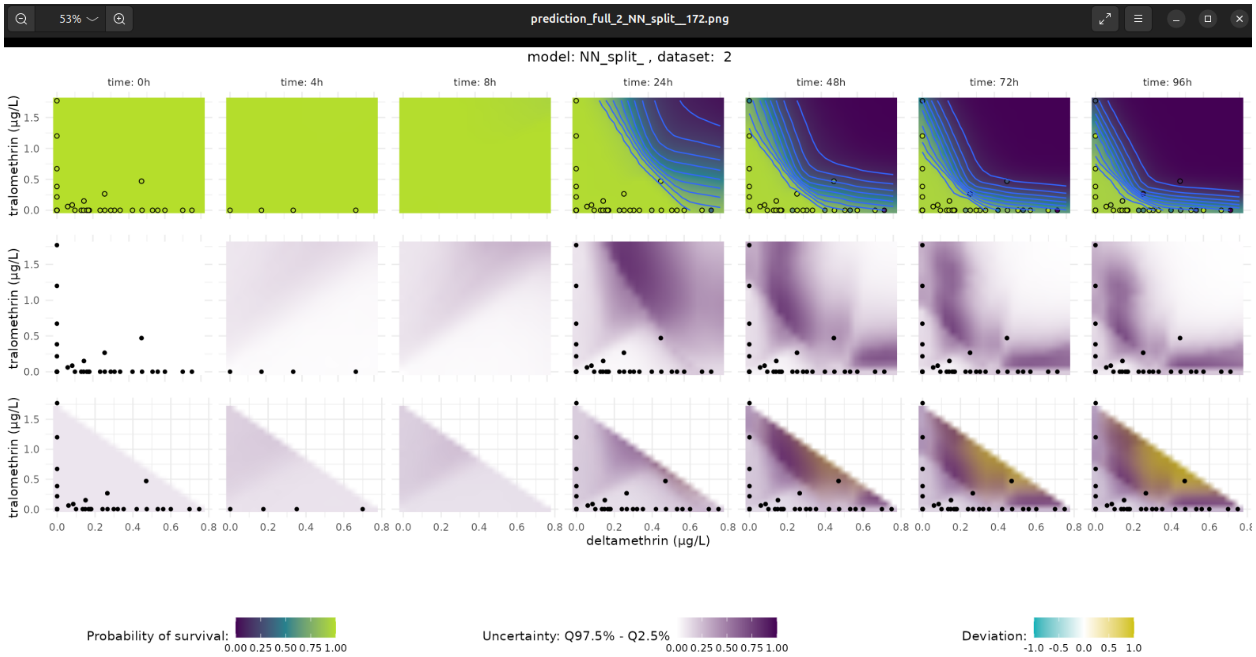
53%
prediction_full_2_NN_split__172.png
model: NN_split_ , dataset: 2
time: 0h
time: 4h
time: 8h
time: 24h
time: 48h
time: 72h
time: 96h
tralomethrin (µg/L)
deltamethrin (µg/L)
Probability of survival:
0.00 0.25 0.50 0.75 1.00
Uncertainty: Q97.5% - Q2.5%
0.00 0.25 0.50 0.75 1.00
Deviation:
-1.0 -0.5 0.0 0.5 1.0

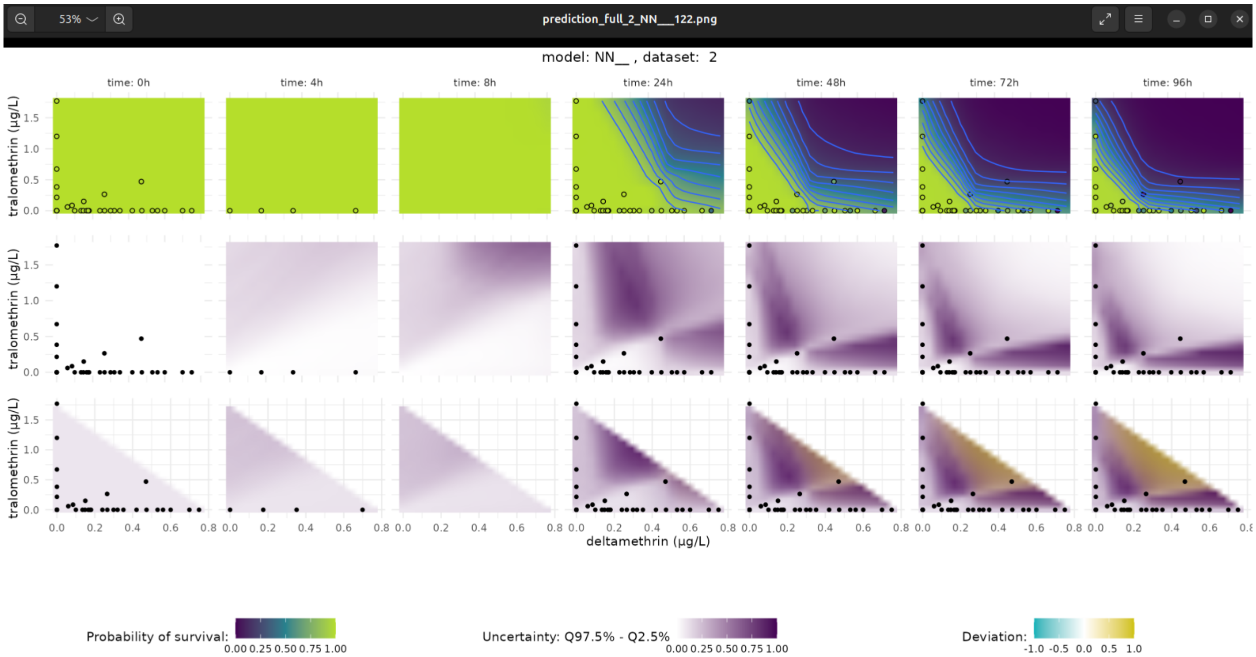
prediction_full_2_NN__122.png
53%
model: NN__ , dataset: 2
time: 0h
time: 4h
time: 8h
time: 24h
time: 48h
time: 72h
time: 96h
tralomethrin (µg/L)
deltamethrin (µg/L)
Probability of survival:
0.00 0.25 0.50 0.75 1.00
Uncertainty: Q97.5% - Q2.5%
0.00 0.25 0.50 0.75 1.00
Deviation:
-1.0 -0.5 0.0 0.5 1.0

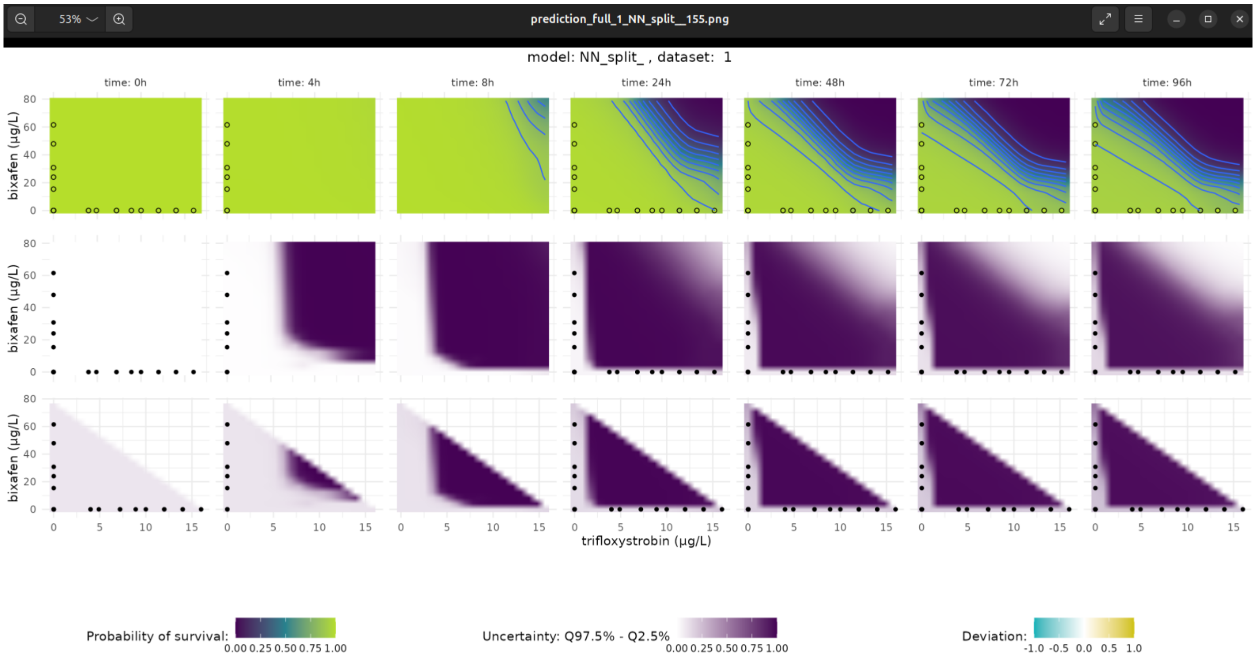
prediction_full_1_NN_split__155.png
53%
model: NN_split_ , dataset: 1
time: 0h
time: 4h
time: 8h
time: 24h
time: 48h
time: 72h
time: 96h
bixafen (µg/L)
trifloxystrobin (µg/L)
Probability of survival:
0.00 0.25 0.50 0.75 1.00
Uncertainty: Q97.5% - Q2.5%
0.00 0.25 0.50 0.75 1.00
Deviation:
-1.0 -0.5 0.0 0.5 1.0

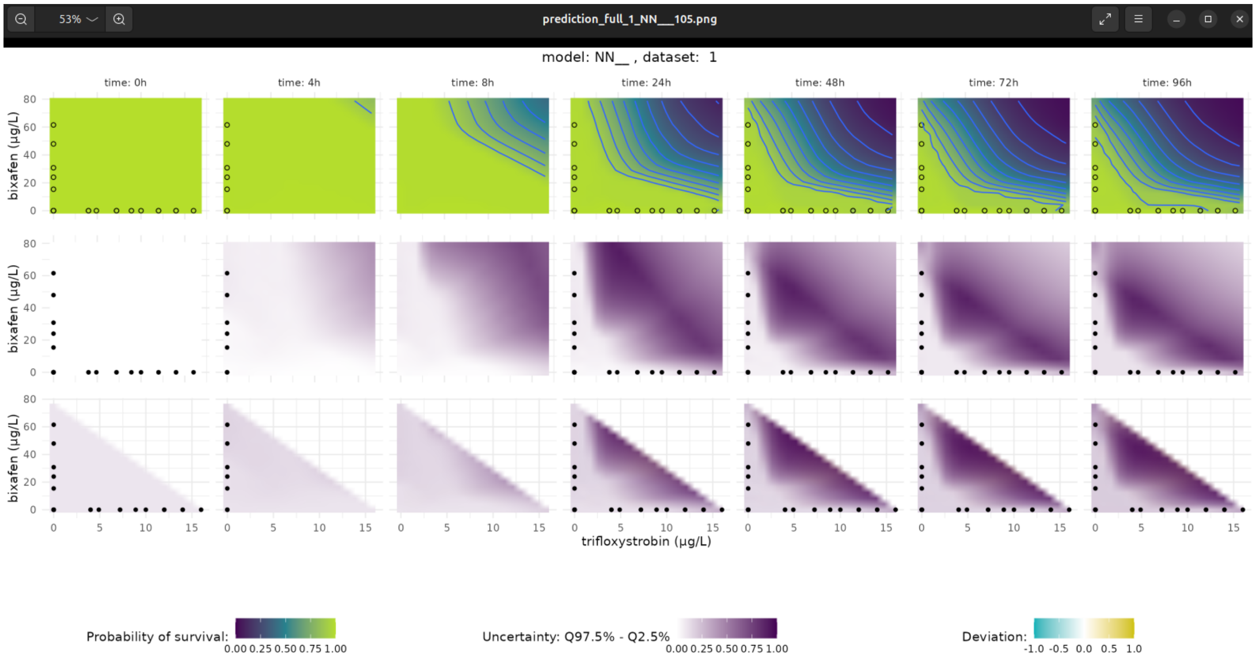
prediction_full_1_NN__105.png
model: NN__ , dataset: 1
time: 0h
time: 4h
time: 8h
time: 24h
time: 48h
time: 72h
time: 96h
bixafen (µg/L)
trifloxystrobin (µg/L)
Probability of survival:
0.00 0.25 0.50 0.75 1.00
Uncertainty: Q97.5% - Q2.5%
0.00 0.25 0.50 0.75 1.00
Deviation:
-1.0 -0.5 0.0 0.5 1.0

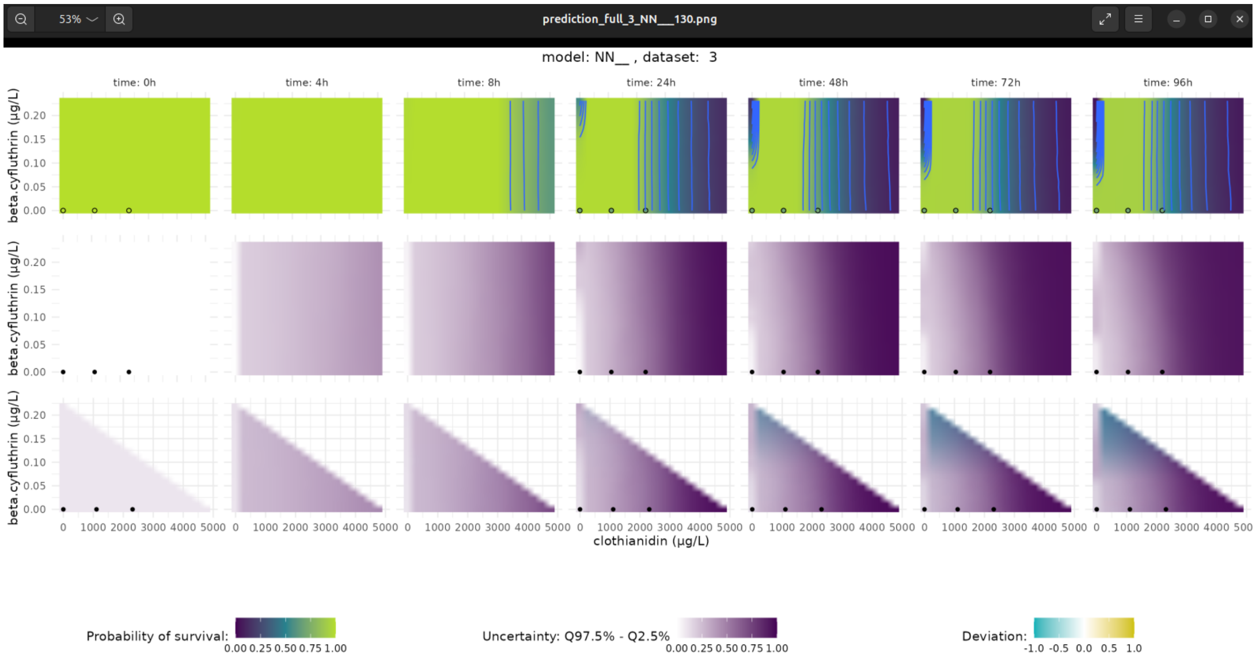
prediction_full_3_NN__130.png
model: NN__ , dataset: 3
time: 0h
time: 4h
time: 8h
time: 24h
time: 48h
time: 72h
time: 96h
beta.cyfluthrin (µg/L)
clothianidin (µg/L)
Probability of survival:
0.00 0.25 0.50 0.75 1.00
Uncertainty: Q97.5% - Q2.5%
0.00 0.25 0.50 0.75 1.00
Deviation:
-1.0 -0.5 0.0 0.5 1.0

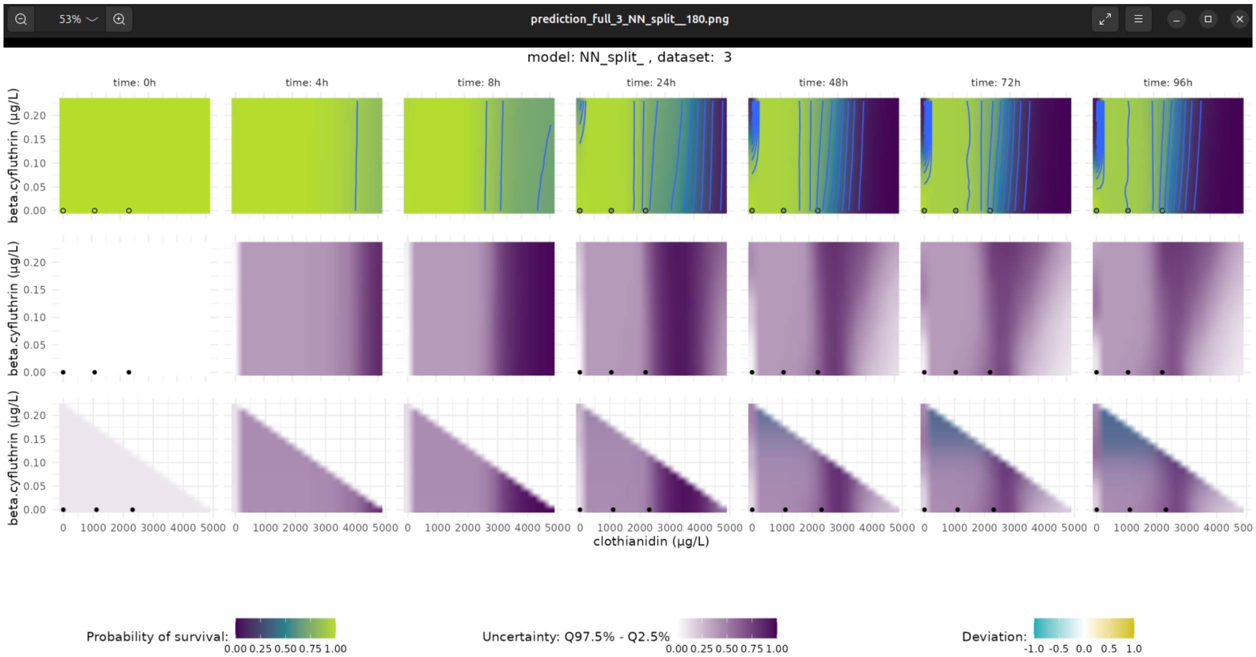
53%
prediction_full_3_NN_split__180.png
model: NN_split_ , dataset: 3
time: 0h
time: 4h
time: 8h
time: 24h
time: 48h
time: 72h
time: 96h
beta.cyfluthrin (µg/L)
0.00 0.05 0.10 0.15 0.20
0 1000 2000 3000 4000 5000
clothianidin (µg/L)
Probability of survival:
0.00 0.25 0.50 0.75 1.00
Uncertainty: Q97.5% - Q2.5%
0.00 0.25 0.50 0.75 1.00
Deviation:
-1.0 -0.5 0.0 0.5 1.0

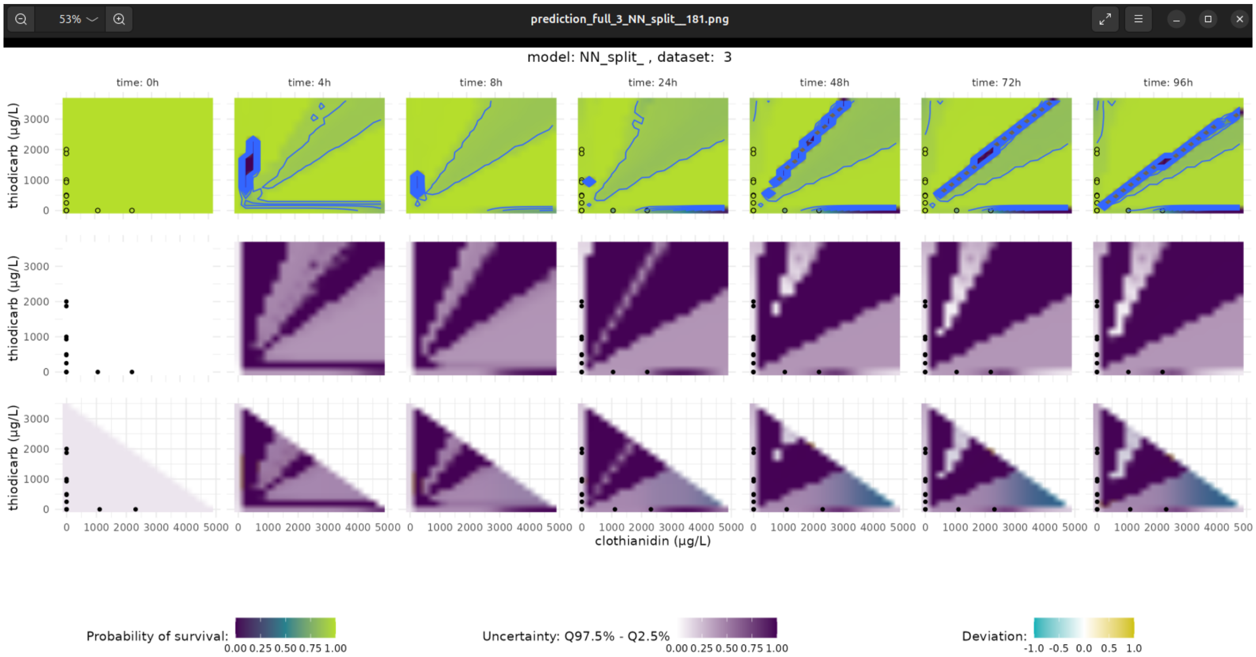
53%
prediction_full_3_NN_split__181.png
model: NN_split_ , dataset: 3
time: 0h
time: 4h
time: 8h
time: 24h
time: 48h
time: 72h
time: 96h
thiodicarb (µg/L)
0 1000 2000 3000
0 1000 2000 3000 4000 5000
clothianidin (µg/L)
Probability of survival:
0.00 0.25 0.50 0.75 1.00
Uncertainty: Q97.5% - Q2.5%
0.00 0.25 0.50 0.75 1.00
Deviation:
-1.0 -0.5 0.0 0.5 1.0

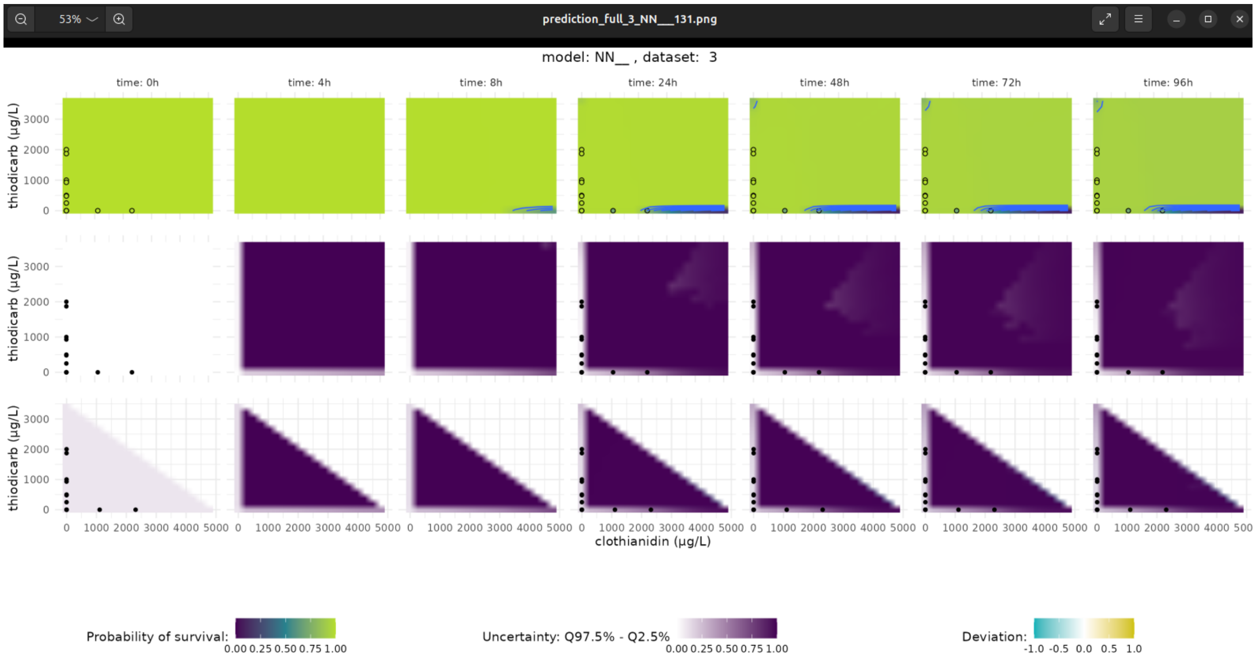
prediction_full_3_NN__131.png
model: NN__ , dataset: 3
time: 0h
time: 4h
time: 8h
time: 24h
time: 48h
time: 72h
time: 96h
thiodicarb (µg/L)
clothianidin (µg/L)
Probability of survival:
0.00 0.25 0.50 0.75 1.00
Uncertainty: Q97.5% - Q2.5%
0.00 0.25 0.50 0.75 1.00
Deviation:
-1.0 -0.5 0.0 0.5 1.0

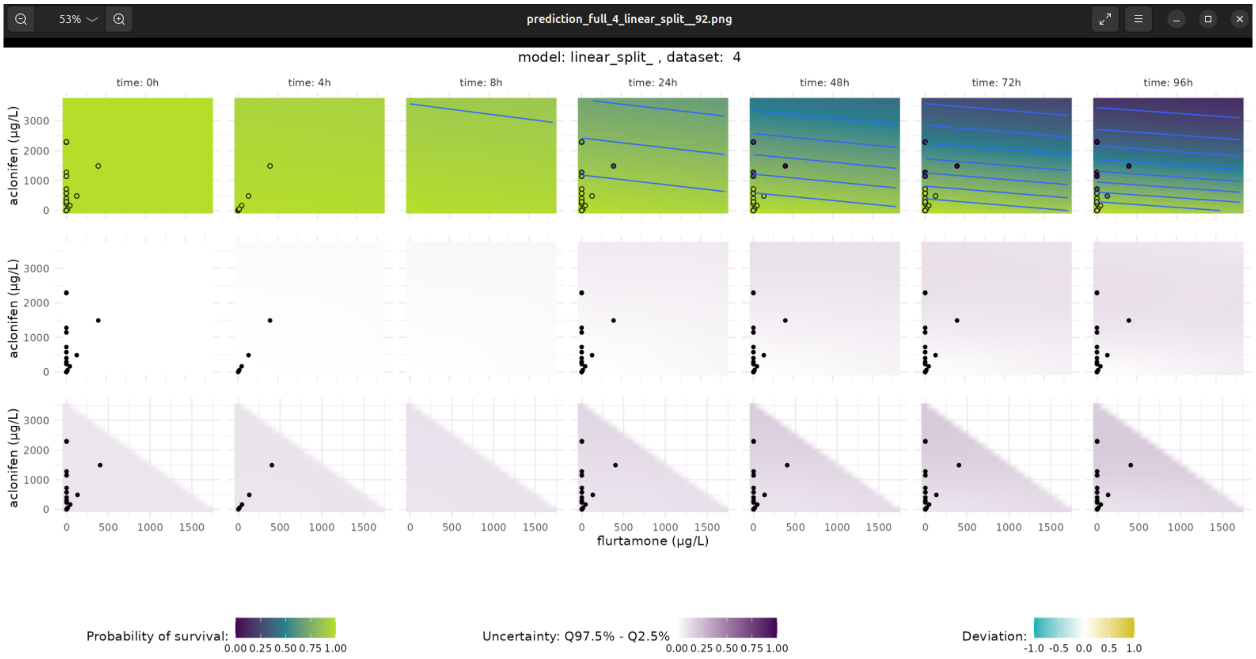
prediction_full_4_linear_split__92.png
model: linear_split_ , dataset: 4
time: 0h
time: 4h
time: 8h
time: 24h
time: 48h
time: 72h
time: 96h
aclonifen (µg/L)
flurtamone (µg/L)
Probability of survival:
0.00 0.25 0.50 0.75 1.00
Uncertainty: Q97.5% - Q2.5%
0.00 0.25 0.50 0.75 1.00
Deviation:
-1.0 -0.5 0.0 0.5 1.0

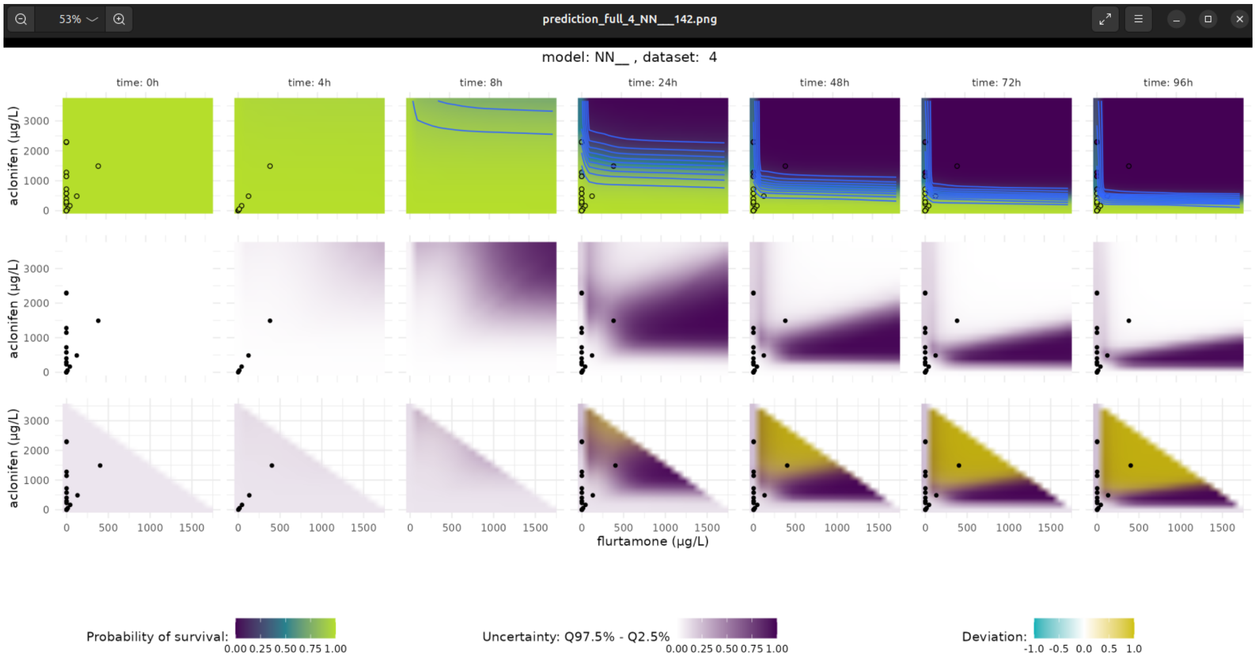
prediction_full_4_NN__142.png
model: NN__ , dataset: 4
time: 0h
time: 4h
time: 8h
time: 24h
time: 48h
time: 72h
time: 96h
aclonifen (µg/L)
flurtamone (µg/L)
Probability of survival:
0.00 0.25 0.50 0.75 1.00
Uncertainty: Q97.5% - Q2.5%
0.00 0.25 0.50 0.75 1.00
Deviation:
-1.0 -0.5 0.0 0.5 1.0

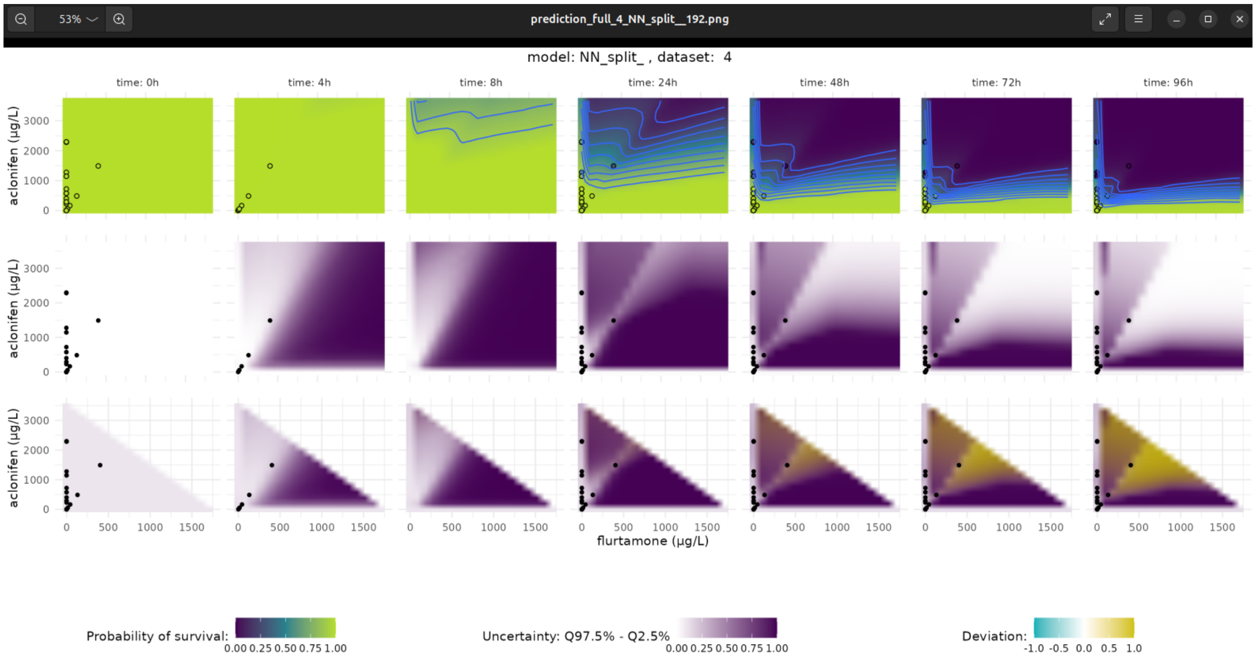
prediction_full_4_NN_split__192.png
model: NN_split_ , dataset: 4
time: 0h
time: 4h
time: 8h
time: 24h
time: 48h
time: 72h
time: 96h
aclonifen (µg/L)
flurtamone (µg/L)
Probability of survival:
0.00 0.25 0.50 0.75 1.00
Uncertainty: Q97.5% - Q2.5%
0.00 0.25 0.50 0.75 1.00
Deviation:
-1.0 -0.5 0.0 0.5 1.0

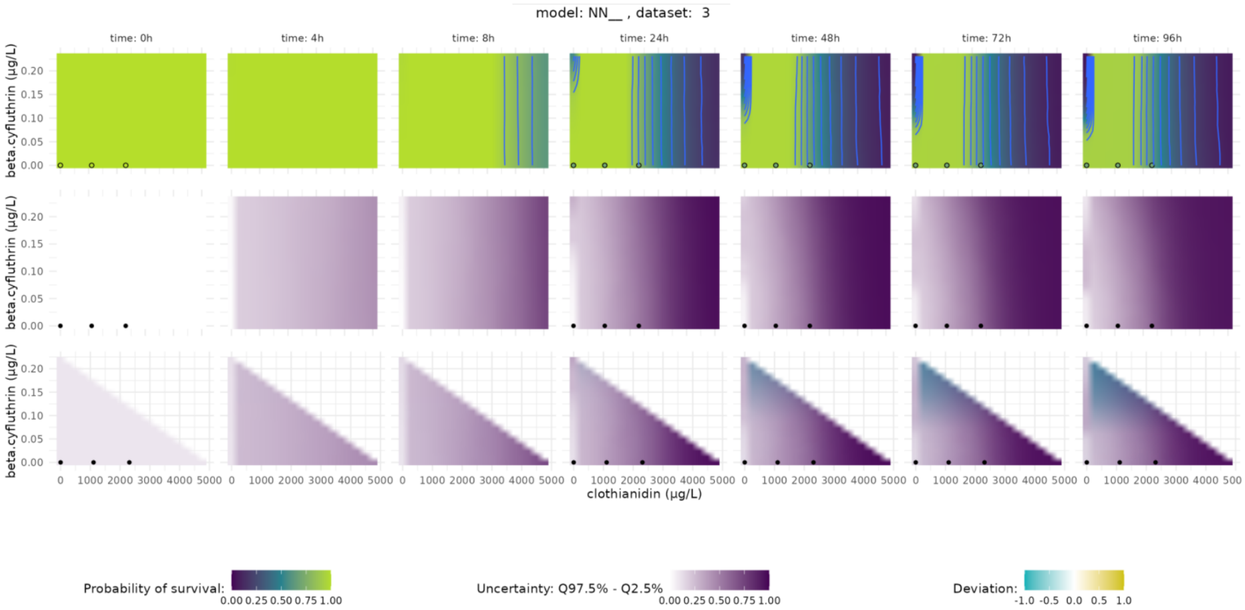
model: NN__ , dataset: 3
time: 0h
time: 4h
time: 8h
time: 24h
time: 48h
time: 72h
time: 96h
beta.cyfluthrin (µg/L)
0.00 0.05 0.10 0.15 0.20
0 1000 2000 3000 4000 5000
clothianidin (µg/L)
Probability of survival:
0.00 0.25 0.50 0.75 1.00
Uncertainty: Q97.5% - Q2.5%
0.00 0.25 0.50 0.75 1.00
Deviation:
-1.0 -0.5 0.0 0.5 1.0

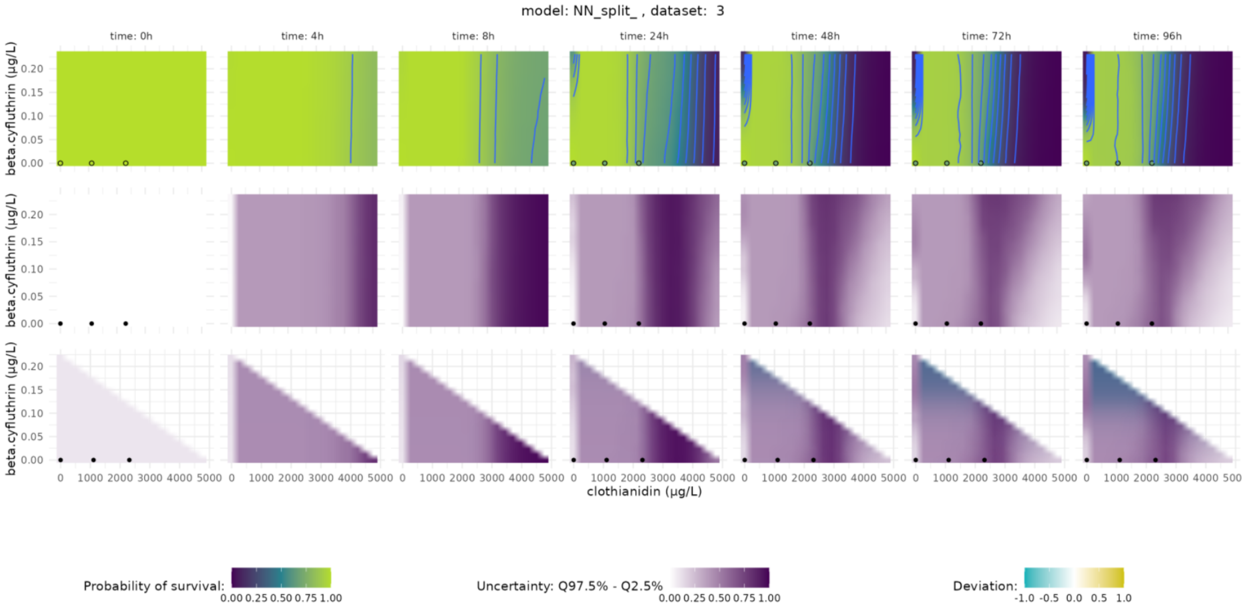
model: NN_split_ , dataset: 3
time: 0h
time: 4h
time: 8h
time: 24h
time: 48h
time: 72h
time: 96h
beta.cyfluthrin (µg/L)
0.00 0.05 0.10 0.15 0.20
0 1000 2000 3000 4000 5000
clothianidin (µg/L)
Probability of survival:
0.00 0.25 0.50 0.75 1.00
Uncertainty: Q97.5% - Q2.5%
0.00 0.25 0.50 0.75 1.00
Deviation:
-1.0 -0.5 0.0 0.5 1.0

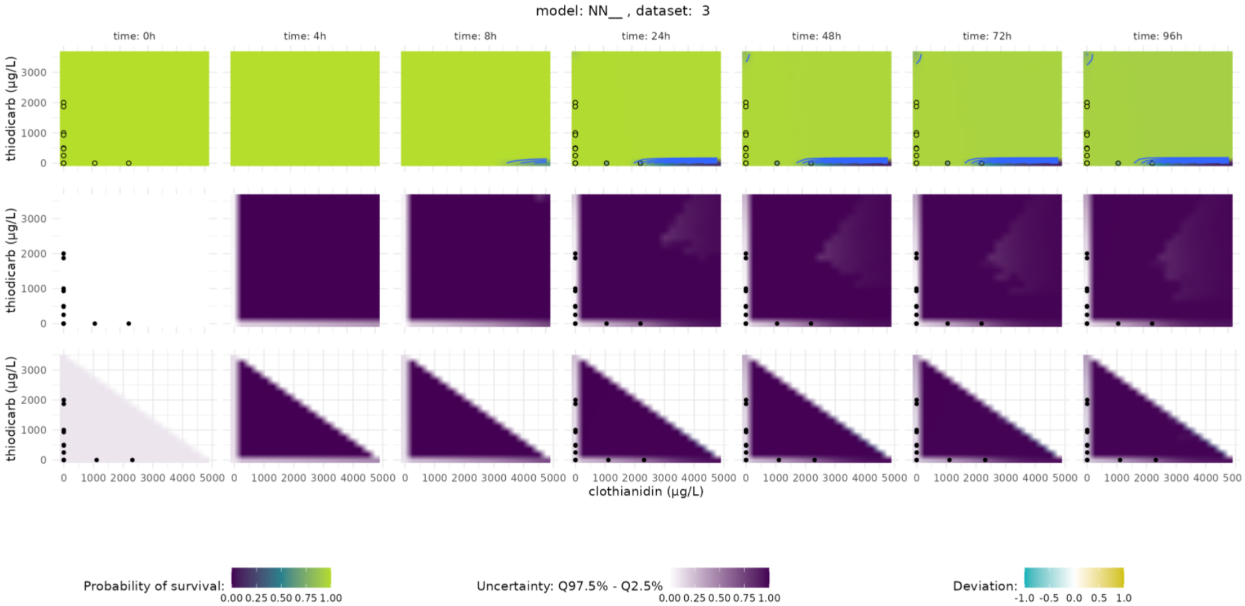
model: NN__ , dataset: 3
time: 0h
time: 4h
time: 8h
time: 24h
time: 48h
time: 72h
time: 96h
thiodicarb (μg/L)
clothianidin (μg/L)
Probability of survival:
0.00 0.25 0.50 0.75 1.00
Uncertainty: Q97.5% - Q2.5%
0.00 0.25 0.50 0.75 1.00
Deviation:
-1.0 -0.5 0.0 0.5 1.0

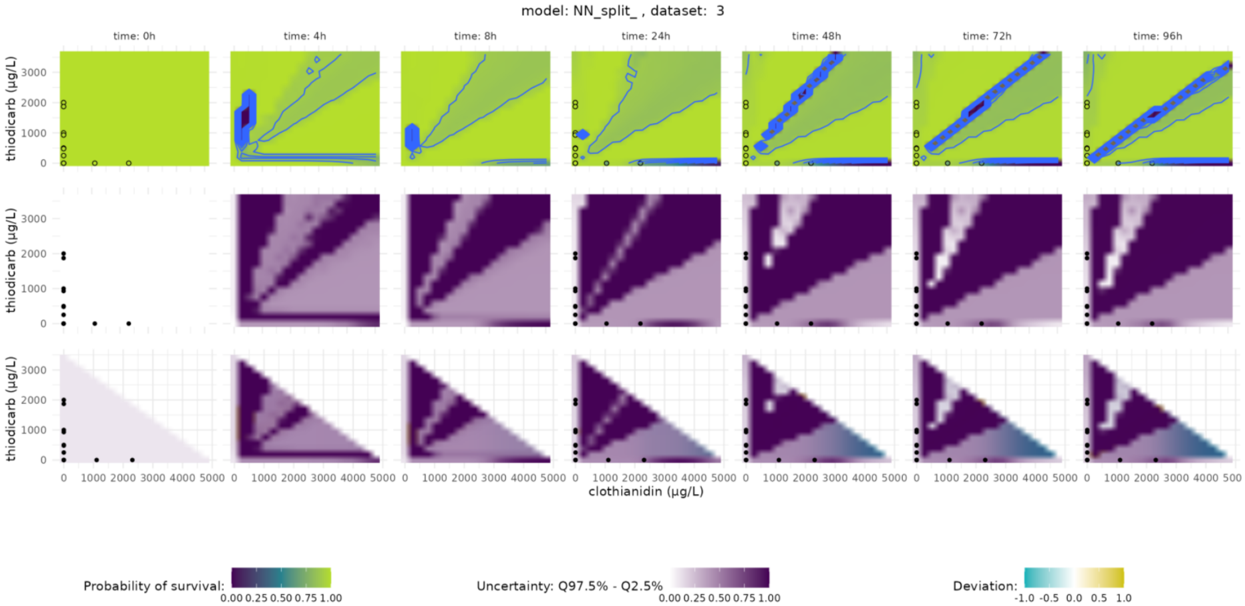
model: NN_split_ , dataset: 3
time: 0h
time: 4h
time: 8h
time: 24h
time: 48h
time: 72h
time: 96h
thiodicarb (μg/L)
clothianidin (μg/L)
Probability of survival:
0.00 0.25 0.50 0.75 1.00
Uncertainty: Q97.5% - Q2.5%
0.00 0.25 0.50 0.75 1.00
Deviation:
-1.0 -0.5 0.0 0.5 1.0

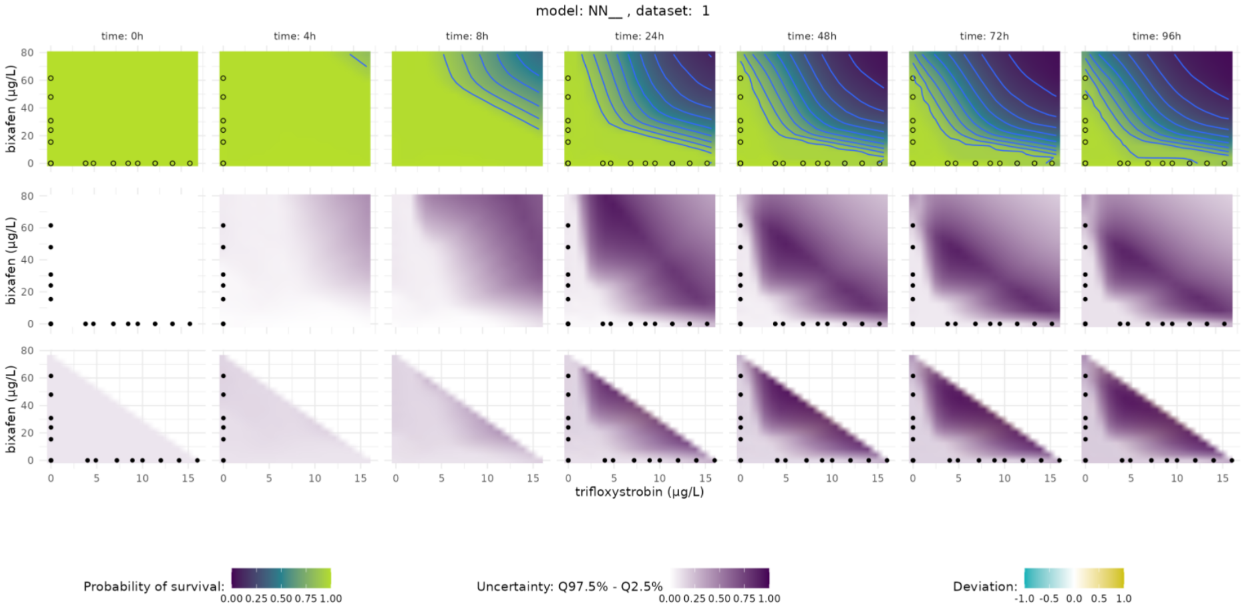
model: NN__ , dataset: 1
time: 0h
time: 4h
time: 8h
time: 24h
time: 48h
time: 72h
time: 96h
bixafen (µg/L)
trifloxystrobin (µg/L)
Probability of survival:
0.00 0.25 0.50 0.75 1.00
Uncertainty: Q97.5% - Q2.5%
0.00 0.25 0.50 0.75 1.00
Deviation:
-1.0 -0.5 0.0 0.5 1.0

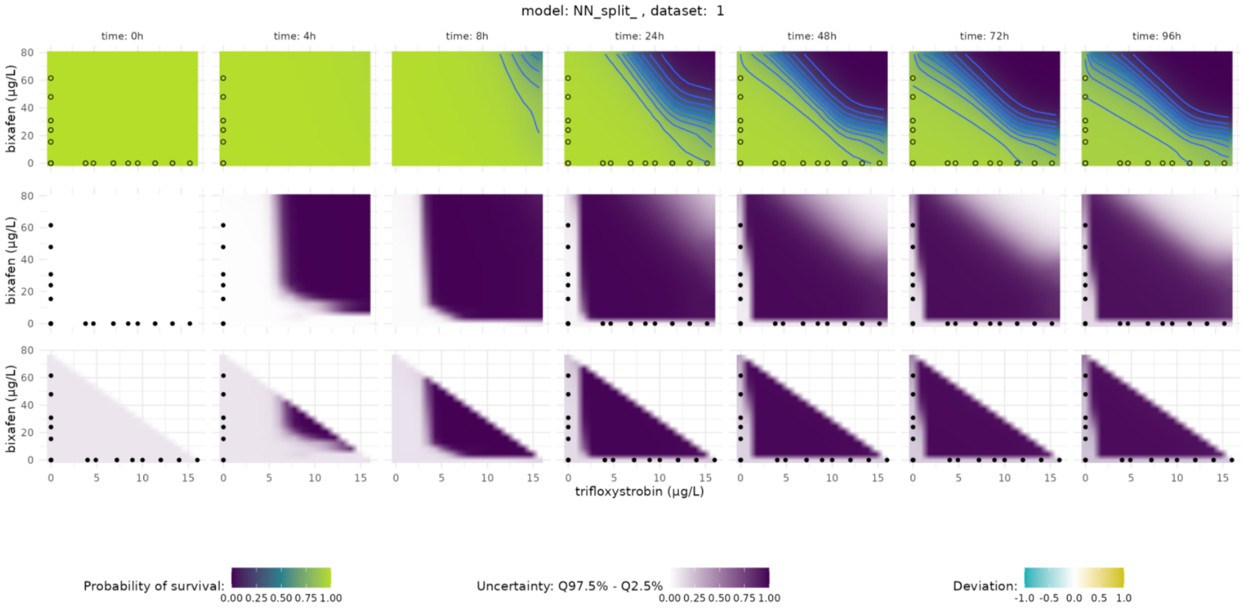
model: NN_split_ , dataset: 1
time: 0h
time: 4h
time: 8h
time: 24h
time: 48h
time: 72h
time: 96h
bixafen (µg/L)
trifloxystrobin (µg/L)
Probability of survival:
0.00 0.25 0.50 0.75 1.00
Uncertainty: Q97.5% - Q2.5%
0.00 0.25 0.50 0.75 1.00
Deviation:
-1.0 -0.5 0.0 0.5 1.0

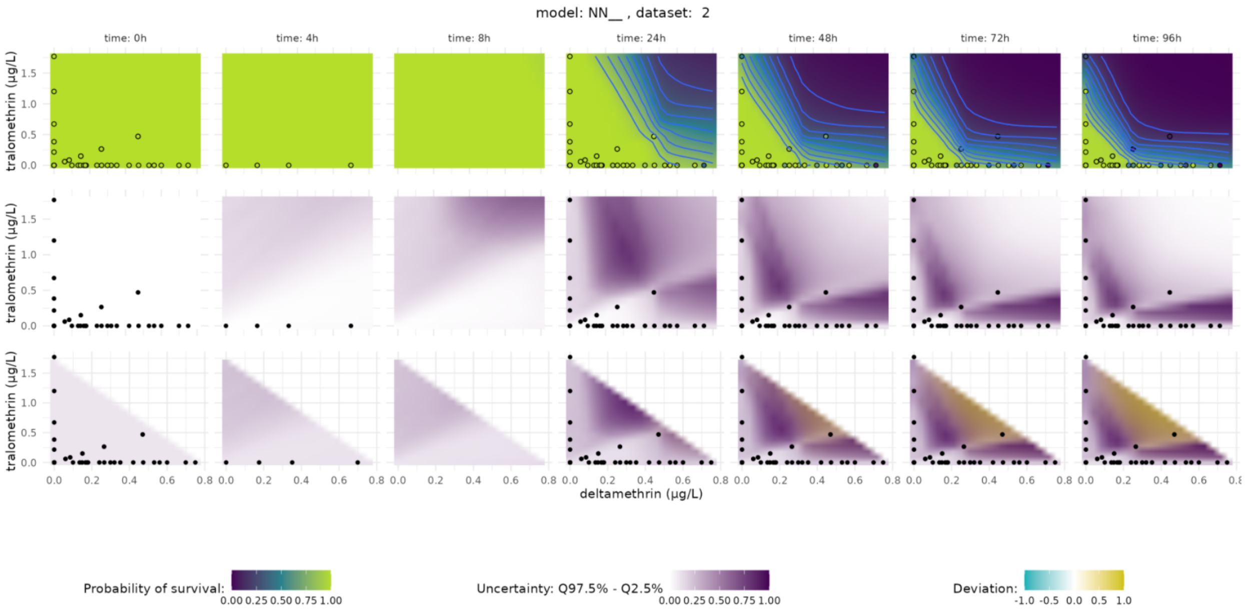
model: NN__ , dataset: 2
time: 0h
time: 4h
time: 8h
time: 24h
time: 48h
time: 72h
time: 96h
tralomethrin (µg/L)
deltamethrin (µg/L)
Probability of survival:
0.00 0.25 0.50 0.75 1.00
Uncertainty: Q97.5% - Q2.5%
0.00 0.25 0.50 0.75 1.00
Deviation:
-1.0 -0.5 0.0 0.5 1.0

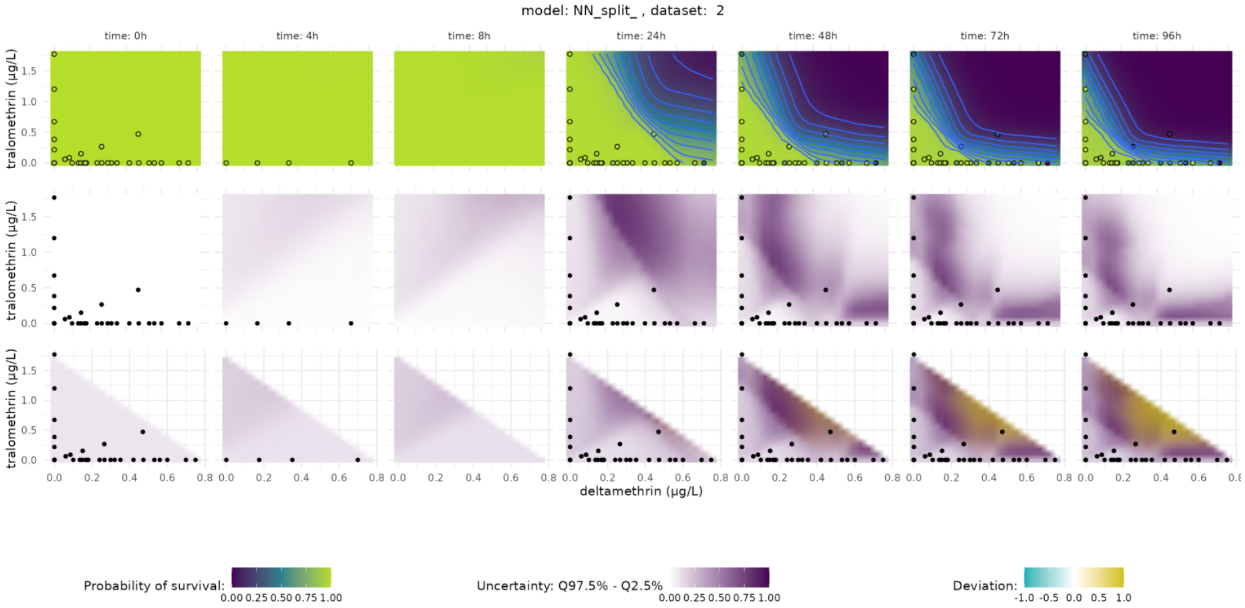
model: NN_split_ , dataset: 2
time: 0h
time: 4h
time: 8h
time: 24h
time: 48h
time: 72h
time: 96h
tralomethrin (µg/L)
deltamethrin (µg/L)
Probability of survival:
0.00 0.25 0.50 0.75 1.00
Uncertainty: Q97.5% - Q2.5%
0.00 0.25 0.50 0.75 1.00
Deviation:
-1.0 -0.5 0.0 0.5 1.0

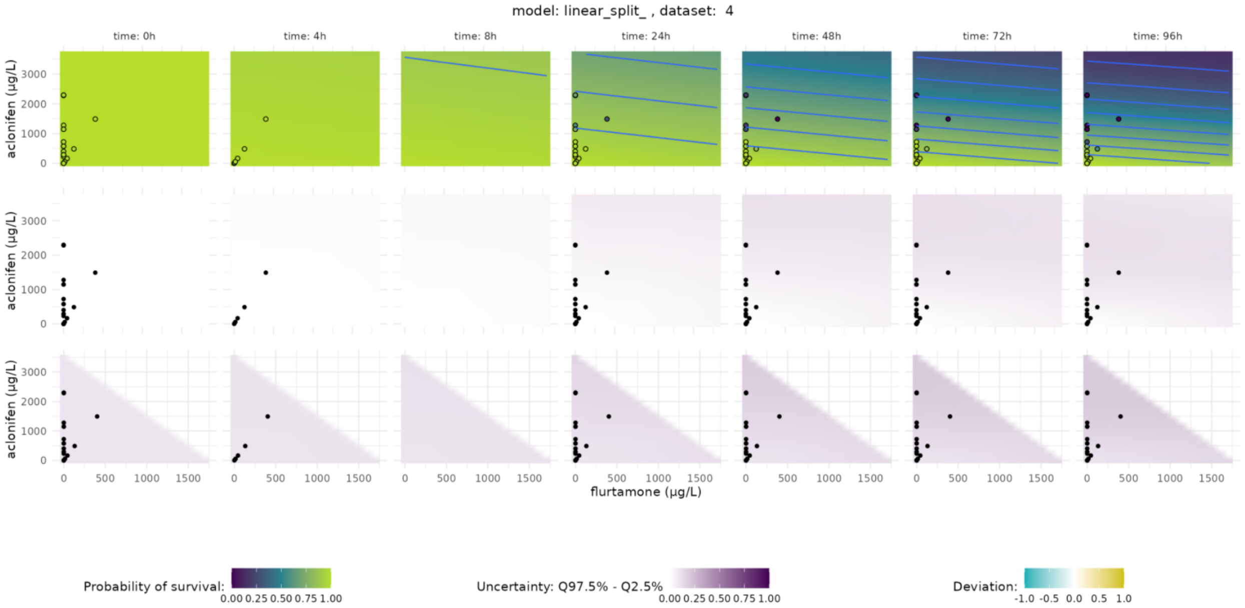
model: linear_split_ , dataset: 4
time: 0h
time: 4h
time: 8h
time: 24h
time: 48h
time: 72h
time: 96h
aclonifen (µg/L)
flurtamone (µg/L)
Probability of survival:
0.00 0.25 0.50 0.75 1.00
Uncertainty: Q97.5% - Q2.5%
0.00 0.25 0.50 0.75 1.00
Deviation:
-1.0 -0.5 0.0 0.5 1.0

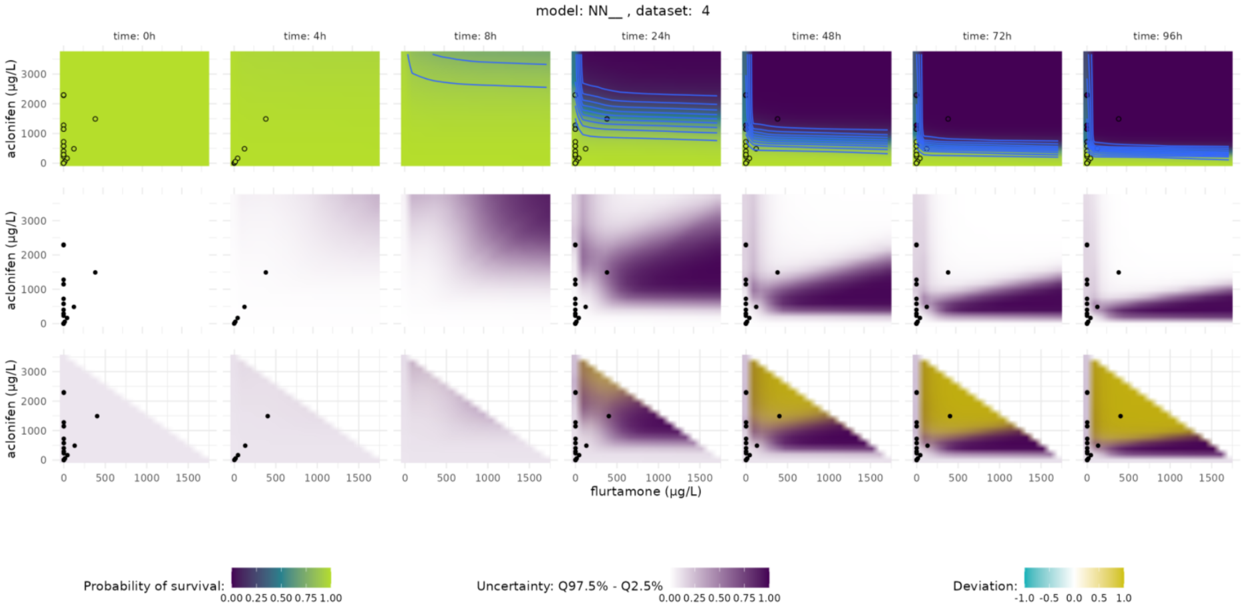
model: NN__ , dataset: 4
time: 0h
time: 4h
time: 8h
time: 24h
time: 48h
time: 72h
time: 96h
aclonifen (µg/L)
flurtamone (µg/L)
Probability of survival:
0.00 0.25 0.50 0.75 1.00
Uncertainty: Q97.5% - Q2.5%
0.00 0.25 0.50 0.75 1.00
Deviation:
-1.0 -0.5 0.0 0.5 1.0

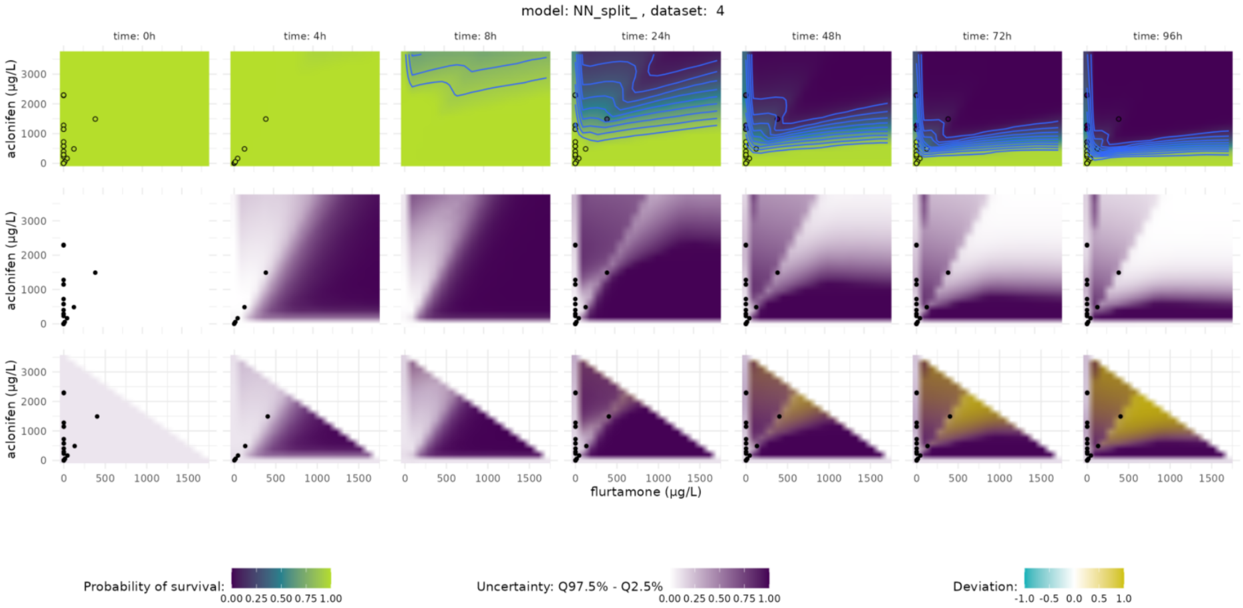
model: NN_split_ , dataset: 4
time: 0h
time: 4h
time: 8h
time: 24h
time: 48h
time: 72h
time: 96h
aclonifen (µg/L)
flurtamone (µg/L)
Probability of survival:
0.00 0.25 0.50 0.75 1.00
Uncertainty: Q97.5% - Q2.5%
0.00 0.25 0.50 0.75 1.00
Deviation:
-1.0 -0.5 0.0 0.5 1.0
