## Supplementary Material - Script and Datasets for "A Bayesian Neural Ordinary Differential Equations Framework to Study the Effects of Chemical Mixtures on Survival": Bayes_TKlinearTD.R

```
library("coda")
library("rjags")

BRIDGE = "linear"
N_UPDATE = 5000
N_ITER = 2500
N_ITER_WAIC = 1500
############################
uniq_rep = unique(d$replicate)
ls_Nprec = lapply(uniq_rep, function(r){
    dat = d[d$replicate == r,]
    a = dat$Nsurv
    Nprec = c(a[1],a[-length(a)]) 
    return(Nprec)
})
Nprec = do.call("c", ls_Nprec)
ls_tprec = lapply(uniq_rep, function(r){
    dat = d[d$replicate == r,]
    a = dat$time
    tprec = c(a[1],a[-length(a)]) 
    return(tprec)
})
tprec = do.call("c", ls_tprec)
d$i_row = 1:nrow(d)
d$lag_i_row = c(d$i_row [1],d$i_row [-length(d$i_row )]) 
d$i_prec = ifelse(d$time == 0, d$i_row, d$lag_i_row)
iprec = d$i_prec
ls_replicate_ID = lapply(seq_along(uniq_rep), function(i){
    dat = d[d$replicate == (uniq_rep[i]),]
    a = rep(i, length(dat$replicate) )
    return(a)
})
replicate_ID = do.call("c", ls_replicate_ID)
ls_time_ID = lapply(uniq_rep, function(r){
    dat = d[d$replicate == r,]
    a = 1:length(dat$time) 
    return(a)
})
time_ID = do.call("c", ls_time_ID)

########## DATA FOR JAGS
m_layer = N_EXPOSURE # hidden layer size
data_IT = list(
    n_data = nrow(d),
    time = d$time,
    time_ID = time_ID,
    replicate_ID = replicate_ID,
    i_prec = iprec,
    Nsurv = d$Nsurv,
    Nprec = Nprec,
    n_X = N_EXPOSURE,
    X = matrix(MIXTURE, ncol = N_EXPOSURE, byrow = FALSE),
    kd_meanlog10 = rep(-1.5,N_EXPOSURE),
    kd_sdlog10 = rep(0.5,N_EXPOSURE),
    hb_meanlog10 = -1.5,
    hb_sdlog10 = 0.5,
    alpha_meanlog10 = -1.5,
    alpha_sdlog10 = 0.5,
    beta_minlog10 = -2,
    beta_maxlog10 = 2,
    hb_value = 1,
    hb_valueFIXED = 0,
    # Neural Network
    m_layer = m_layer
)

### Model building
print(paste("START MODEL", ID_SET, BRIDGE))
model <- jags.model(
        file = "src/JAGS_TKlinearTD_IT.txt",
        data = data_IT,
        n.chains = 3
    )

print(paste("START UPDATE", ID_SET, BRIDGE))
update(model, N_UPDATE)

print(paste("START TRACE", ID_SET, BRIDGE))
mcmc_trace <- coda.samples(
    model,
    c("a", "b1", "kd", "hb", "alpha", "beta", "Nsurv_ppc", "Nsurv_sim"),
    n.iter = N_ITER)

print(paste("START WAIC", ID_SET, BRIDGE))
load.module("dic")
mcmc_mean <- jags.samples(
    model, c("deviance", "WAIC"),
    type = "mean",
    n.iter = N_ITER_WAIC)
# sapply(mcmc_mean, sum)

path = paste0("output/mcmc_", ID_SET, "_JAGS_TK",BRIDGE,"TD_IT.rda")
save(file = path, model, mcmc_mean, mcmc_trace, d)

############ PLOT
#print(paste("START PLOT", ID_SET, BRIDGE))
#extract_chain = function(mcmc, key){
#    ls_mcmc = lapply(mcmc, function(m){
#        matching_columns = grepl(key, colnames(m))
#        return(m[,matching_columns])
#    })
#    matrix_key = do.call("rbind",ls_mcmc)
#    return(matrix_key)
#}
#matrix_mcmc = extract_chain(mcmc_trace, 'Nsurv_sim')
#prediction = apply(matrix_mcmc, 2, quantile, c(0.05,0.5,0.975))

#d$med = prediction[2,]
#d$qinf = prediction[1,]
#d$qsup = prediction[3,]
#plt = ggplot(data = d) + 
#    theme_minimal() +
#    geom_point(aes(x = time, y = Nsurv)) +
#    geom_ribbon(aes(x = time, ymin = qinf, ymax = qsup), color = "lightgrey", alpha = 0.5) +
#    geom_line(aes(x = time, y = med), color = "red") +
#    facet_wrap(~ replicate)
#path_png = paste0("output/mcmc_", ID_SET, "_JAGS_TK",BRIDGE,"TD_IT.png")
#ggsave(path_png, plt)

####### COPY FILE
# system("cp -r output/. ~/../mnt/output", intern = TRUE)
```
