## Supplementary Material - Script and Datasets for "A Bayesian Neural Ordinary Differential Equations Framework to Study the Effects of Chemical Mixtures on Survival": README.md

# Maxim TK NeuralODEs TD

# Using neural ordinary differential equations to predict chemical mixture effects on survival

This is the companion repository for the paper *[Using neural ordinary differential equations to predict chemical mixture effects on survival](/paper.pdf)* by [Virgile Baudrot](https://juanmc2005.github.io/), Nina Cedergreen, Thomas Kleiber, André Gergs and Sandrine Charles.

> Plant Protection Products (PPP) are formulated to maximise their efficacy to control target pest species, including mixtures of active substances. 
Non-target species are likely exposed to mixtures of PPP due to their combined use within PPP formulations. More significantly, these species may encounter untested mixtures resulting from different uses, as PPP may be applied at different locations and timings in the landscape. PPP thus undergo various degradation processes, resulting in potentially countless mixture exposure profiles.
Being able to predict joint effects of active substances without having to test all combinations in the laboratory increases the efficiency of risk assessment models.
As toxicity is a process that occurs over time, evaluating the effects over time gives valuable information on the toxicity of both single chemicals and mixtures.
Time-variable effects can be assessed with toxicokinetic (TK) and toxicodynamic (TD) models. The mixture toxicity concepts (Concentration Addition and Independent Action) have recently been implemented in survival TKTD models, as means to deviate from the models through interactions with the limiting rate constant of the TKTD models.
However, this approach cannot account for the nonlinear mechanisms of synergies and antagonisms, whether at the level of internal concentrations (TK) or observed effects (TD).
Here we provide a standalone pipeline coupling TKTD mechanistic models with neural networks (NN) to map potential interactions between active substances in combinations or in formulations without prior expectations in terms of mechanisms by which interactions may occur.
Using this approach benefits from both the mechanistic and NN worlds to fit experimental data (under a Bayesian framework) on active substances alone and in combination with oher active substances or additives.
Ultimately, this helps identify chemical mixtures that deviate from the reference models of Concentration Addition and Independent Action and predict interactions of new combinations and formulations not yet tested.
Our model has been tested based on 99 acute toxicity studies in which fish survival has been quantified, grouped into 4 data sets, with 5 to 6 different active substances used in varying concentrations and formulations. 
The study presents a first analysis that demonstrates the robustness of the hybridization of the mechanistic model of Ordinary Differential Equation (ODE) with Neural Networks (NN) and the inference approach, together with preliminary results on the type of interaction that seems to occur most often.

## Install

1) Create R environment with JAGS and the library `Rjags`:

```shell
sudo apt-get update
sudo apt-get -y install r-base
apt-get -y update && apt-get -y install jags
R -e "install.packages('coda',dependencies=TRUE, repos='http://cran.rstudio.com/')"
R -e "install.packages('rjags',dependencies=TRUE, repos='http://cran.rstudio.com/')"
```

2) Copy all require files:

The variable `MIP` is the `IP` of the remote machine.

```shell
MIP="XXX.XXX.XXX.XXX"
ssh root@$MIP

mkdir src
mkdir data
mkdir output
```

Copy JAGS models:

```shell
scp -r src/JAGS_TKTD_IT_n.txt root@$MIP:~/src/
scp -r src/JAGS_TKTD_IT_n_exp.txt root@$MIP:~/src/
scp -r src/JAGS_TKTD_IT_nn_n_exp.txt root@$MIP:~/src/
scp -r src/JAGS_TKTD_IT_nn_ReLU_n_exp.txt root@$MIP:~/src/
scp -r src/JAGS_TKTD_IT_nn_ReLU_nn_ReLU_n_exp.txt root@$MIP:~/src/
scp -r src/JAGS_TKTD_IT_nn_ReLU_nn_ReLU_nn_ReLU_n_exp.txt root@$MIP:~/src/
```

Copy R scripts:

```shell
scp -r Bayes_TKTD_n.R root@$MIP:~/
scp -r Bayes_TKTD_n_exp.R root@$MIP:~/
scp -r Bayes_TKTD_nn_n_exp.R root@$MIP:~/
scp -r Bayes_TKTD_nn_ReLU_n_exp.R root@$MIP:~/
scp -r Bayes_TKTD_nn_ReLU_nn_ReLU_n_exp.R root@$MIP:~/
scp -r Bayes_TKTD_nn_ReLU_nn_ReLU_nn_ReLU_n_exp.R root@$MIP:~/
```

Copy Data:

```shell
scp -r data/data_artificial_synergism.csv root@$MIP:~/data/
scp -r data/data_artificial_antagonism.csv root@$MIP:~/data/
scp -r data/data_artificial_additive.csv root@$MIP:~/data/

scp -r data/MaXim__raw_datasets__set1_CLEAN.csv root@$MIP:~/data/
scp -r data/MaXim__raw_datasets__set2_CLEAN.csv root@$MIP:~/data/
scp -r data/MaXim__raw_datasets__set3_CLEAN.csv root@$MIP:~/data/
scp -r data/MaXim__raw_datasets__set4_CLEAN.csv root@$MIP:~/data/
```

Copy run files:

```shell
scp -r run_*.R root@$mip:~/
```

## Run an experiment

- Use `run_*.R` to reproduce the results.


## Citation

```bibtex
@article{baudrot2024TKTDNeural,
  author={Baudrot, Virgile and Cedergreen, Nina and Kleiber, Thomas and Gergs, André and Charles, Sandrine},
  title={{Using neural ordinary differential equations to predict chemical mixture effects on survival}}, 
  year={2024},
  volume={},
  number={},
  pages={},
  doi={}
}
```

## License

```
MIT License

Copyright (c) 2024 Qonfluens

Permission is hereby granted, free of charge, to any person obtaining a copy
of this software and associated documentation files (the "Software"), to deal
in the Software without restriction, including without limitation the rights
to use, copy, modify, merge, publish, distribute, sublicense, and/or sell
copies of the Software, and to permit persons to whom the Software is
furnished to do so, subject to the following conditions:

The above copyright notice and this permission notice shall be included in all
copies or substantial portions of the Software.

THE SOFTWARE IS PROVIDED "AS IS", WITHOUT WARRANTY OF ANY KIND, EXPRESS OR
IMPLIED, INCLUDING BUT NOT LIMITED TO THE WARRANTIES OF MERCHANTABILITY,
FITNESS FOR A PARTICULAR PURPOSE AND NONINFRINGEMENT. IN NO EVENT SHALL THE
AUTHORS OR COPYRIGHT HOLDERS BE LIABLE FOR ANY CLAIM, DAMAGES OR OTHER
LIABILITY, WHETHER IN AN ACTION OF CONTRACT, TORT OR OTHERWISE, ARISING FROM,
OUT OF OR IN CONNECTION WITH THE SOFTWARE OR THE USE OR OTHER DEALINGS IN THE
SOFTWARE.
```
