## Supplementary Material - Script and Datasets for "A Bayesian Neural Ordinary Differential Equations Framework to Study the Effects of Chemical Mixtures on Survival": run_all2.R

```
library("coda")
library("rjags")

# 2
ID_SET = "2"
d = read.delim("data/MaXim__raw_datasets__set2_CLEAN.csv", header = TRUE, sep = ",") 
colnames(d)
MIXTURE = c(d$spiromesifen, d$deltamethrin, d$triazophos, d$tralomethrin, d$flupyradifurone)
N_EXPOSURE = 5
source("Bayes_TKTD_n.R")
source("Bayes_TKTD_n_exp.R")
source("Bayes_TKTD_nn_ReLU_n_exp.R")
source("Bayes_TKTD_nn_ReLU_nn_ReLU_n_exp.R")
source("Bayes_TKTD_nn_ReLU_nn_ReLU_nn_ReLU_n_exp.R")
```
