## Supplementary Material - Script and Datasets for "A Bayesian Neural Ordinary Differential Equations Framework to Study the Effects of Chemical Mixtures on Survival": run_all4.R

```
library("coda")
library("rjags")

# 4
ID_SET = "4"
d = read.delim("data/MaXim__raw_datasets__set4_CLEAN.csv", header = TRUE, sep = ",") 
colnames(d)
MIXTURE = c(d$flufenacet, d$diflufenican, d$metribuzin, d$flurtamone, d$aclonifen)
N_EXPOSURE = 5
source("Bayes_TKTD_n.R")
source("Bayes_TKTD_n_exp.R")
source("Bayes_TKTD_nn_ReLU_n_exp.R")
source("Bayes_TKTD_nn_ReLU_nn_ReLU_n_exp.R")
source("Bayes_TKTD_nn_ReLU_nn_ReLU_nn_ReLU_n_exp.R")
```
