## Supplementary Material - Script and Datasets for "A Bayesian Neural Ordinary Differential Equations Framework to Study the Effects of Chemical Mixtures on Survival": run_all_ant.R

```
library("coda")
library("rjags")

# antagonism
ID_SET = "antagonism"
d = read.delim("data/data_artificial_antagonism.csv", header = TRUE, sep = ",") 
colnames(d)
MIXTURE = c(d$A, d$B)
N_EXPOSURE = 2
source("Bayes_TKTD_n.R")
source("Bayes_TKTD_n_exp.R")
source("Bayes_TKTD_nn_ReLU_n_exp.R")
source("Bayes_TKTD_nn_ReLU_nn_ReLU_n_exp.R")
source("Bayes_TKTD_nn_ReLU_nn_ReLU_nn_ReLU_n_exp.R")
```
