## Supplementary Material - Script and Datasets for "A Bayesian Neural Ordinary Differential Equations Framework to Study the Effects of Chemical Mixtures on Survival": JAGS_TKTD_IT_nn_ReLU_nn_ReLU_n_exp.txt

model {
    ##------------------------------------------------------ priors
    ## -- TOXICOKINETICS - TK
    for (i in 1:n_X){
        kd_taulog10[i] <- 1 / kd_sdlog10[i]^2
        kd_log10[i] ~ dnorm(kd_meanlog10[i], kd_taulog10[i])
        ### -- parameter transformation
        kd[i] <- 10**kd_log10[i]
    }
    # -- TK-TD BRIDGE
    for(i in 1:m_layer){
        for(j in 1:m_layer){
            W1[i,j] ~ dnorm(0.0, 4)
        }
        b1[i] ~ dnorm(0.0, 4)
    }
    for(i in 1:m_layer){
        for( j in 1:m_layer){
            W2[i,j] ~ dnorm(0.0, 4)
        }
        b2[i] ~ dnorm(0.0, 4) 
    }
    for(i in 1:1){
        for( j in 1:m_layer){
            W3[i,j] ~ dnorm(0.0, 4)
        }
        b3 ~ dnorm(0.0, 4) 
    }
  ## -- TOXICODYNAMICS - TD
  hb_taulog10 <- 1 / hb_sdlog10^2
  alpha_taulog10 <- 1 / alpha_sdlog10^2

  hb_log10 ~ dnorm(hb_meanlog10, hb_taulog10)
  alpha_log10 ~ dnorm(alpha_meanlog10, alpha_taulog10)
  beta_log10 ~ dunif(beta_minlog10, beta_maxlog10) # unif -2, 2

  ##---------------------------------------- parameter transformation
  hb <- ifelse(hb_value == 0, hb_valueFIXED, 10**hb_log10)
  alpha <- 10**alpha_log10
  beta <- 10**beta_log10

  ##------------------------------------ model
  for( i in 1:n_data){
    for(j in 1:n_X){
        D_[i,j] <- X[i,j] * ( 1 - exp( - kd[j] * time[i] ))
    }
    nn_D1[i,1:m_layer] = W1 %*% D_[i,1:m_layer] + b1
    for(j in 1:m_layer){
        a_D1[i,j] = max(nn_D1[i,j], 0)
    }
    nn_D2[i,1:m_layer] = W2 %*% a_D1[i,1:m_layer] + b2
    for(j in 1:m_layer){
        a_D2[i,j] = max(nn_D2[i,j], 0)
    }
    nn_D3[i] = W3 %*% a_D2[i,1:m_layer] + b3
    # we exp_transform because we need D > 0
    D[replicate_ID[i], time_ID[i]] <- exp(nn_D3[i])

    D_max[replicate_ID[i], time_ID[i]] <- max(D[replicate_ID[i], 1:time_ID[i]])
    F[i]  <- D_max[replicate_ID[i], time_ID[i]]**beta / ( D_max[replicate_ID[i], time_ID[i]]**beta + alpha**beta )

    psurv_pred[i] <-  exp(-hb * time[i]) * (1 - F[i])
    
    # positivity psurv[i] for division
    psurv[i] <- ifelse(psurv_pred[i] > 0, psurv_pred[i], 1e-10)
    # Ensure 0 < ratio psurv < 1 (JAGS does not support p=1!)
    ratio_pred[i] <- psurv[i]/psurv[i_prec[i]]
    ratio[i] <- ifelse(ratio_pred[i] < 1, 
                        ifelse( ratio_pred[i] > 0,  ratio_pred[i], 1e-10),
                      1-1e-10)    
    Nsurv[i] ~ dbin(ratio[i] , Nprec[i])

 ## ---------------------- generated data 
    Nsurv_ppc[i] ~ dbin(ratio[i] , Nprec[i])
  }
  ## initialization is requires to use 'Nsurv_sim[i-1]' require in JAGS language (avoid auto-loop issue).

    Nsurv_sim[1] ~ dbin(ratio[1], Nprec[1])
    for( i in 2:n_data){
        Nsurv_sim[i] ~ dbin(ratio[i], ifelse( i == i_prec[i], Nprec[i], Nsurv_sim[i-1]))
    }
}
    
